# Ethylene Receptor Gain- and Loss-of-function Mutants Reveal an ETR1-dependent Transcriptional Network in Roots

**DOI:** 10.1101/2024.06.26.600793

**Authors:** Maleana G. White, Alexandria F. Harkey, Joëlle K. Mühlemann, Amy L. Olex, Nathan J. Pfeffer, Maarten Houben, Brad M. Binder, Gloria K. Muday

## Abstract

In Arabidopsis, a family of five receptors mediates ethylene responses in roots, with Ethylene Response 1 (ETR1) controlling increases in root hair proliferation and decreases in lateral root formation. To define the ETR1-dependent gene regulatory network (GRN) controlling root development, we profiled the root transcriptome from Col-0 and the *etr1-3* gain-of-function and *etr1-7* loss-of-function mutants in the presence and absence of ethylene or the ethylene precursor 1-aminocyclopropane-1-carboxylic acid (ACC). We identified 4,522 differentially expressed (DE) transcripts in Col-0 roots with altered abundance in response to ethylene and/or ACC treatment, with larger magnitude changes induced by ethylene. These included 553 DE transcripts that were ETR1 dependent, defined by a lack of response to treatment with ethylene and/or ACC in ethylene-insensitive *etr1-3* and constitutively altered in *etr1-7* in the presence and absence of treatment relative to time-0 Col-0. Within these ETR1-dependent transcripts were ethylene biosynthesis genes and transcription factors. *ACC OXIDASE 2* (*ACO2)* and *ACO3* convert ACC to ethylene and were ETR1-dependent, and *ACO*-promoter-driven reporter fusions were ACC regulated in root tissues in appropriate locations to control root development, with *ACO5* localized to root hairs. Abundance of ETR1-dependent transcripts that were predicted to encode transcription factors and ACOs were examined in Col-0 and an *ein3eil1* mutant with and without ACC treatment, suggesting the ETR1 and EIN3/EIL1 canonical ethylene signaling pathway regulated some, but not all, of these transcriptional responses. Together, these findings reveal features of an ETR1-dependent GRN that controls both ethylene synthesis and root growth and development.

**One sentence summary:** Transcriptional responses in *etr1* LOF and GOF mutants reveal an ethylene-mediated ETR1- and EIN3 dependent gene regulatory network that modulates ethylene signaling and synthesis and root development.

## Introduction

Ethylene is a gaseous hormone that modulates growth and development throughout a plant’s life cycle, from seed germination to senescence, and controls responses to both biotic and abiotic stress (Abeles et al., 1992; Van de Poel et al., 2015). In young, etiolated seedlings, ethylene induces a triple response: reduced hypocotyl and root elongation, thickening of the hypocotyl, and exaggeration of the apical hook. This triple response phenotype has been the basis of mutant screens in *Arabidopsis thaliana* that have identified key components of the ethylene signaling pathway with ethylene-insensitive or constitutive ethylene signaling phenotypes (Azhar et al., 2020; Schaller and Kieber, 2002). Ethylene also has profound effects on root development, including inhibition of primary root elongation and lateral root formation and stimulation of root hair initiation and elongation (Feng et al., 2017; Harkey et al., 2018; Lewis et al., 2011; Negi et al., 2008; Qin et al., 2019). Many of the proteins that were identified based on their functions in ethylene signaling in etiolated hypocotyls also modulate root development (Khoury et al., 2024b), but with distinct developmental effects downstream of each receptor (Harkey et al., 2018).

Ethylene functions through direct binding to members of a family of transmembrane receptors localized to the endoplasmic reticulum and the Golgi apparatus (Azhar et al., 2020; Binder, 2020; Schaller and Kieber, 2002). These receptors are negative regulators of the pathway, such that binding of ethylene to these receptors removes repression of the pathway, which initiates a signaling cascade that upregulates or downregulates genes with distinct transcriptional responses depending on the tissue type and growth conditions (Chang et al., 2013; Harkey et al., 2018, 2019; Stepanova et al., 2007). In Arabidopsis, five receptor isoforms have been identified: Ethylene Response 1 (ETR1), ETR2, Ethylene Insensitive 4 (EIN4), Ethylene Response Sensor 1 (ERS1), and ERS2 (Binder, 2020; Chang et al., 1993; Hua et al., 1998; Hua and Meyerowitz, 1998; Sakai et al., 1998; Schaller and Bleecker, 1995). Unlike most canonical signaling pathways, ethylene receptors are active in the unbound state. In the absence of ethylene, these receptors activate the Raf-like serine/threonine protein kinase Constitutive Triple Response 1 (CTR1), destabilizing EIN3 and EIN3-like (EIL) transcription factors (TFs) and blocking changes in gene expression downstream of these TFs (An et al., 2010; Binder, 2020; Dolgikh et al., 2019; Kieber et al., 1993). In the bound state, however, the ethylene receptors lead to stabilization of EIN3 and EIL TFs in an EIN2-dependent manner (Alonso et al., 1999; An et al., 2010; Huang et al., 2003; Kieber et al., 1993; Qiao et al., 2009), leading to genome-wide changes in transcript abundance (Chang et al., 2013; Harkey et al., 2018). Downstream of EIN3/EIL are ethylene response factors (ERFs) and ethylene response DNA-binding factors (EDFs) that further mediate transcription, consistent with the existence of tissue-specific gene regulatory networks (GRNs) (Wang et al., 2002).

The function of ethylene signaling proteins in controlling root architecture has been examined (Khoury et al., 2024b). EIN2 is required for root developmental responses to ethylene and its immediate precursor 1-aminocyclopropane-1-carboxylic acid (ACC) at 1 µM (Alonso and Stepanova, 2004; Negi et al., 2008), although a recent report indicates that treatment with ACC at levels between 10 and 40 µM can alter root development in *ein2-5* (Mou et al., 2025). The *ctr1-1* mutant was demonstrated to have constitutive-ethylene-signaling root phenotypes, with this mutant having shorter primary roots and increased root hair proliferation compared to Col-0 seedlings in the absence of ethylene (Hua and Meyerowitz, 1998; Kieber et al., 1993; Masucci and Schiefelbein, 1996; Rahman et al., 2000), suggesting its presence is required to suppress the pathway in the absence of ethylene. EIN3 and EIL1 TFs function in ethylene-dependent root hair formation, with the *ein3eil1* double mutant having reduced ACC-induction of root hair proliferation (Feng et al., 2017). However, root developmental responses to ethylene can still be observed in the *ein3* and *ein3eil1* mutants (Chang et al., 2013; Harkey et al., 2018), suggesting that additional TFs may participate in these transcriptional responses to ethylene.

Recent studies have provided new insight into the mechanisms by which ACC is converted to ethylene and identified conditions in which ACC affects plant development independent of its conversion to ethylene. ACC is synthesized by the enzyme ACC synthase from S-adenosyl methionine and then converted to ethylene via ACC oxidase (Houben and Van de Poel, 2019). There are five ACC oxidase enzymes in Arabidopsis, and the transcripts encoding these enzymes have been found to be regulated by ACC and ethylene, thereby increasing ethylene levels in a feed-forward loop. Several quintuple mutants with defects in the five *ACO* genes have now been used to demonstrate that ACC can affect root development in the absence of ACO enzyme activity, providing evidence for actions of ACC independent of its conversion to ethylene (Houben et al., 2024; Mou et al., 2025). The receptors that mediate the regulation of ethylene synthesis by ACC and ethylene, and whether ACC might regulate ACO in an ethylene-independent manner, have not been previously reported.

The five ethylene receptors in Arabidopsis each elicit distinct ethylene responses (Shakeel et al., 2013), partially due to differences in receptor subunit structure and receptor dimerization (Berleth et al., 2019). ETR1 was found to be the main receptor controlling inhibition of ethylene perception by silver nitrate, nutational bending of hypocotyls, and remodeling of root architecture (Harkey et al., 2018; McDaniel and Binder, 2012). In Arabidopsis, ETR1 positively regulates nutational bending (Binder et al., 2006), while the other four receptors negatively regulate this process (Kim et al., 2011). In roots, using a series of single and multiple loss-of-function (LOF) and gain-of-function (GOF) mutants, only ETR1 was needed for the inhibition of lateral root formation by ACC (Harkey et al., 2018). The inhibition of lateral root formation by ACC was lost in the GOF, ethylene-insensitive mutant *etr1-3,* which had significantly more lateral roots than wild type in the presence and absence of ACC. The LOF, constitutive-signaling mutant *etr1-7* had significantly fewer lateral roots than wild type in the presence and absence of ACC. ETR1 also has a central role in the inhibition of primary root elongation and stimulation of root hair formation by ACC, consistent with ETR1 playing the most substantial role in modulating ethylene-mediated root remodeling (Harkey et al., 2018). Nevertheless, while it has been demonstrated that elevated levels of ethylene remodel the root transcriptome (Harkey et al., 2018, 2019; Stepanova et al., 2007), the GRN downstream of the ETR1 receptor has not yet been identified in roots.

To identify the ETR1-dependent GRN controlling ethylene- and ACC-regulated root growth and development, we performed an RNA-Seq analysis in roots from light-grown seedlings using Col-0 and ETR1 receptor GOF (*etr1-3*) and LOF (*etr1-7*) mutants. We identified a core set of genes regulated by either or both ethylene and ACC in Col-0, including a subgroup of these transcripts that we defined as ETR1-dependent due to mis-regulation in the mutants. These transcripts had constitutively altered abundance in time-0 *etr1-7* (as compared to time-0 Col-0), and their responses to ACC and/or ethylene were lost in both *etr1-3* and *etr1-7* mutants. We identified ACC- or ethylene-dependent regulation of transcripts encoding the ACOs, which convert ACC to ethylene, revealing a subset that were ETR1-dependent. Analysis of verified ACO transcriptional reporters driven by the native ACO promoters revealed overlapping and distinct root tissue expression patterns among this protein family. Finally, we explored the ETR1-dependent GRN through the lens of ETR1-dependent transcripts predicted to encode TFs, building a network model that revealed TF-encoding transcripts with common temporal and ETR1-regulated responses. We identified those TFs that are targets of EIN3 and tested whether these transcriptional responses required functional copies of EIN3/EIL1. This work sheds light on the ETR1 and EIN3-dependent GRNs that mediate the transcriptional responses to elevated levels of ACC and its ethylene product.

## Results

### Ethylene and ACC doses with similar ETR1-dependent effects on root hair growth

The main goal of this study was to define the genes that depend upon ETR1 for their regulation by ethylene. We examined the transcriptional responses to both ethylene and its immediate precursor ACC to compare their ETR1-dependent GRNs in these experiments, as previous work suggested that ACC and ethylene may act through distinct signaling pathways in some tissues (Houben et al., 2024; Mou et al., 2020; Yin et al., 2019). As a first step, we determined the concentration of ethylene that yielded moderate root hair proliferation like that induced by 0.75 µM ACC (Martin et al., 2022), by examining root hairs in Col-0 seedlings treated with a range of ethylene levels for 24 hours (Supplemental Figure 1). The 0.3 ppm ethylene concentration appeared to induce root hair proliferation to a similar degree as 0.75 µM ACC, so we selected these concentrations for this RNA-Seq analysis. The effects of a 24-hour treatment with these doses of ethylene and ACC were examined in the *etr1-3* and *etr1-7* mutants revealing proliferation of root hairs in the *etr1-7* mutant in the absence of treatment (Figure 1B). The ethylene-insensitive *etr1-3* mutant showed reduced responses to ACC and ethylene, forming fewer and shorter root hairs in the presence of either treatment than Col-0. We used our published ACC time-course microarray dataset (Harkey et al., 2018) to determine the time points that had large numbers of differentially expressed (DE) genes that were not found in other time points and overlapped with the early, middle, and late root hair developmental responses, selecting 1, 4, and 24 hour treatment times (Supplemental Figure 2) (for details, see Methods).

**Figure 1.**
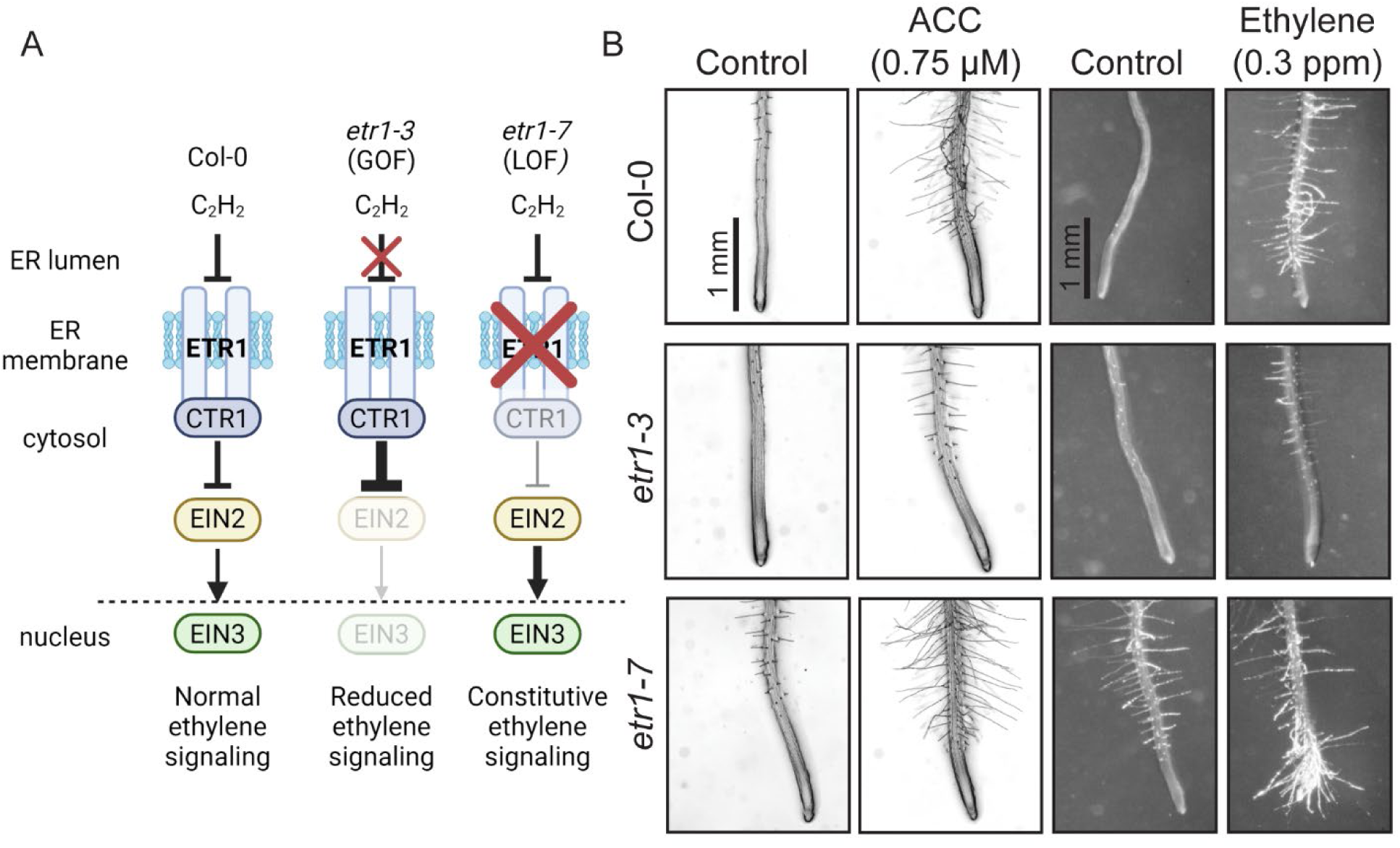
Altered ethylene signaling in the *etr1-3* and *etr1-7* mutants results in aberrant root hair formation. (A) A schematic illustrating the wild-type ethylene signaling pathway and how it is altered in *etr1-3* and *etr1-7* mutants. This image was prepared using Biorender. (B) Representative images of six-day-old Col-0, *etr1-3*, and *etr1-7* seedlings treated with 0.3 ppm ethylene or 0.75 μM ACC for 24 hours, revealing similar developmental responses by these two doses of ethylene and ACC within the different genotypes.

### The reduced effects of ACC and ethylene on transcriptional response in ETR1 mutants are revealed by principal component analysis

We performed this RNA-Seq analysis in triplicate in root samples from three genotypes (Col-0, *etr1-3*, and *etr1-7*), two treatments (0.75 µM ACC and 0.3 ppm ethylene gas), and four time points (0, 1, 4, and 24 hours). After sequencing and quality control, principal component analysis (PCA) plots were created using 1000 genes with the most variance between samples to identify which factors contributed most to this variance. Figure 2A depicts the PCA plot for all samples, which highlights differences between genotypes. The *etr1-7* samples (triangles) are distinct with higher PC1 variance than Col-0 and *etr1-3*. There is a clear separation across PC2 between time-0 samples (in green) and ethylene- and ACC-treated samples (blue, purple, and black symbols). Additionally, one time-0 Col-0 sample separated from other Col-0 time-0 samples along the PC2 axis, which prompted us to analyze variation *within* genotypes. These plots (Supplemental Figure 3; described further in Methods) led us to remove the outlier Col-0 sample from downstream analyses.

**Figure 2.**
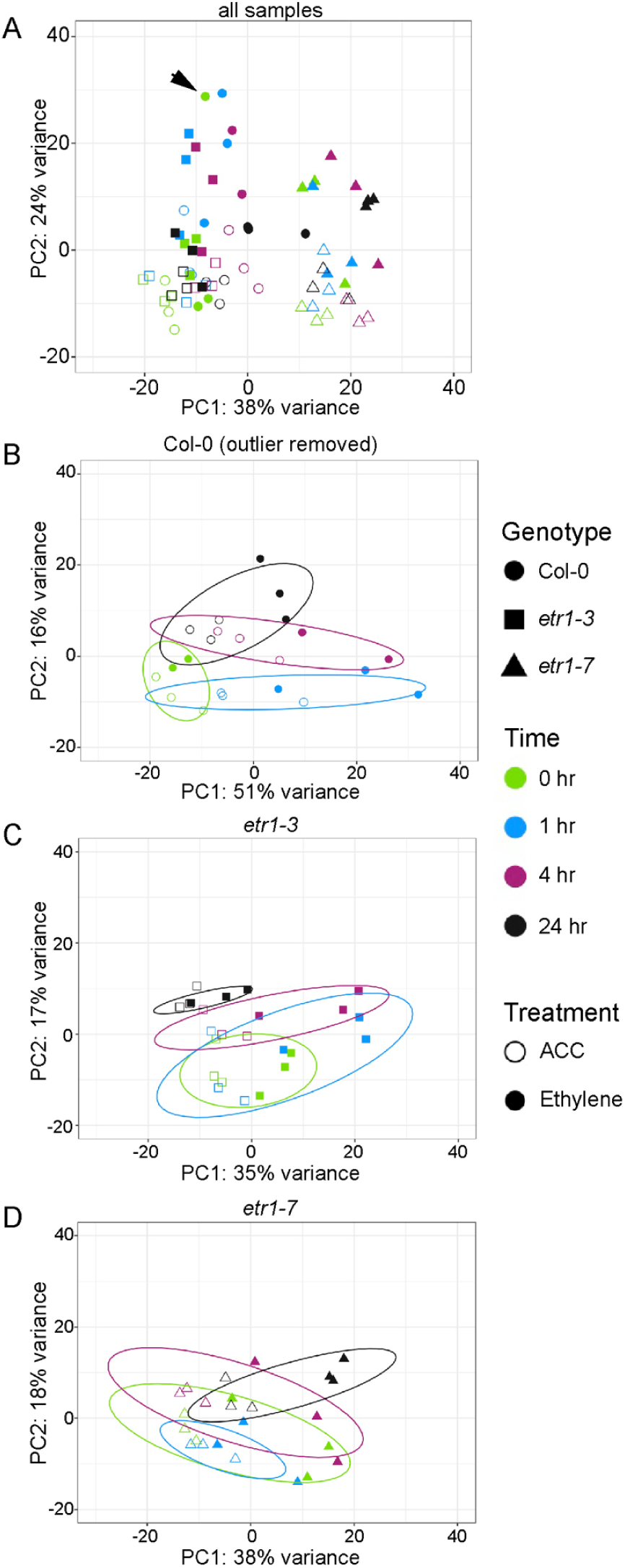
Large-scale transcript abundance patterns in data show *etr1-7* samples are distinctly different from Col-0 and *etr1-3*. (A) PCA plot of all sequenced samples with the genotype indicated by shape of the symbol, the time of treatment indicated by color, and the treatment type ACC or ethylene, indicated by open or filled symbols, respectively. The arrow denotes an untreated Col-0 outlier. PCA plots of (B) Col-0 (with the outlier sample removed), (C) *etr1-3*, and (D) *etr1-7* samples. Circles surround the 80% confidence intervals for each time point.

We used three genotype-specific PCA plots (Figure 2B-D) to assess the contribution of treatment type and duration to variation among samples within genotype. In Col-0 the time 0 samples (green circles) cluster together, consistent with the absence of exposure to ACC or ethylene. The effect of treatment time is most evident in the separation between samples (circled color groups) along the PC2 axes generally shifting from time 0 h to 24 h from the bottom to the top of the axis. This PCA plot also emphasizes the difference between ethylene- and ACC-treated samples across the PC1 axis, explaining 51% of the variance, suggesting unique responses to these two molecules.

An important point that is revealed by these PCA plots is that both *etr1-3* and *etr1-7* mutants have dampened response to treatment with ethylene and ACC, consistent with reduced or enhanced signaling, respectively. In Col-0, PC1, which separates ACC and ethylene treatments in all genotypes, explains 51% of the variance. In the mutants, PC1 explains only 35% and 38% of the variance. Thus, the treatments have smaller effects on overall transcript abundance changes in the mutants. This dampened response is also evident in *etr1-3* by the overlapping responses of time 0 and 1 hour (green and blue samples) and the reduced separation along PC2 for the 4- and 24-hour samples (purple and black), with especially tight clustering in the 24-hour treatment. The *etr-7* samples show even less separation by time of treatment along the PC2 axes (comparison of color of samples), especially for ACC-treated samples (open circles), suggesting that this mutant is exhibiting ethylene and ACC responses in the absence of treatment. Plots of Pearson’s correlation (Supplemental Figure 4A and B) and of Euclidean distance (Supplemental Figure 4C and D) with and without hierarchical clustering support these conclusions, as do manually arranged plots grouped by genotype and time of ACC and ethylene treatment (Supplemental Figure 4A and C; described further in Methods).

### Ethylene treatment stimulated more dramatic transcriptional changes than ACC treatment in Col-0

To summarize how the abundance of transcripts was affected by ethylene and ACC treatment, we first examined the hormone-induced responses in Col-0. For both treatments, every time point was compared back to the same set of combined Col-0 controls, defined above. We created a density plot depicting the number of transcripts as a function of their log_2_FC using those transcripts with a significant adjusted p-value < 0.01 as determined by DESeq2 (Figure 3A). We observed a bimodal distribution at each time point, showing one group of transcripts that increased in abundance and another that decreased in abundance because of these treatments. This pattern was found in response to both ethylene and ACC, although there were larger numbers of DE transcripts and greater magnitude changes in response to ethylene than ACC treatment, especially at one hour. Both treatments resulted in more down-regulated than up- regulated genes at the 4- and 24-hour time points. The bimodal pattern and higher number of down-regulated genes in response to ethylene and ACC at the 4- and 24-hour time points were similar to our previous microarray study using ACC (Harkey et al., 2018). In all treatments and time points, the bimodal peaks representing the increases and decreases in transcript abundance relative to time-0 Col-0 were centered on or near −0.5 and 0.5 log_2_FC.

**Figure 3.**
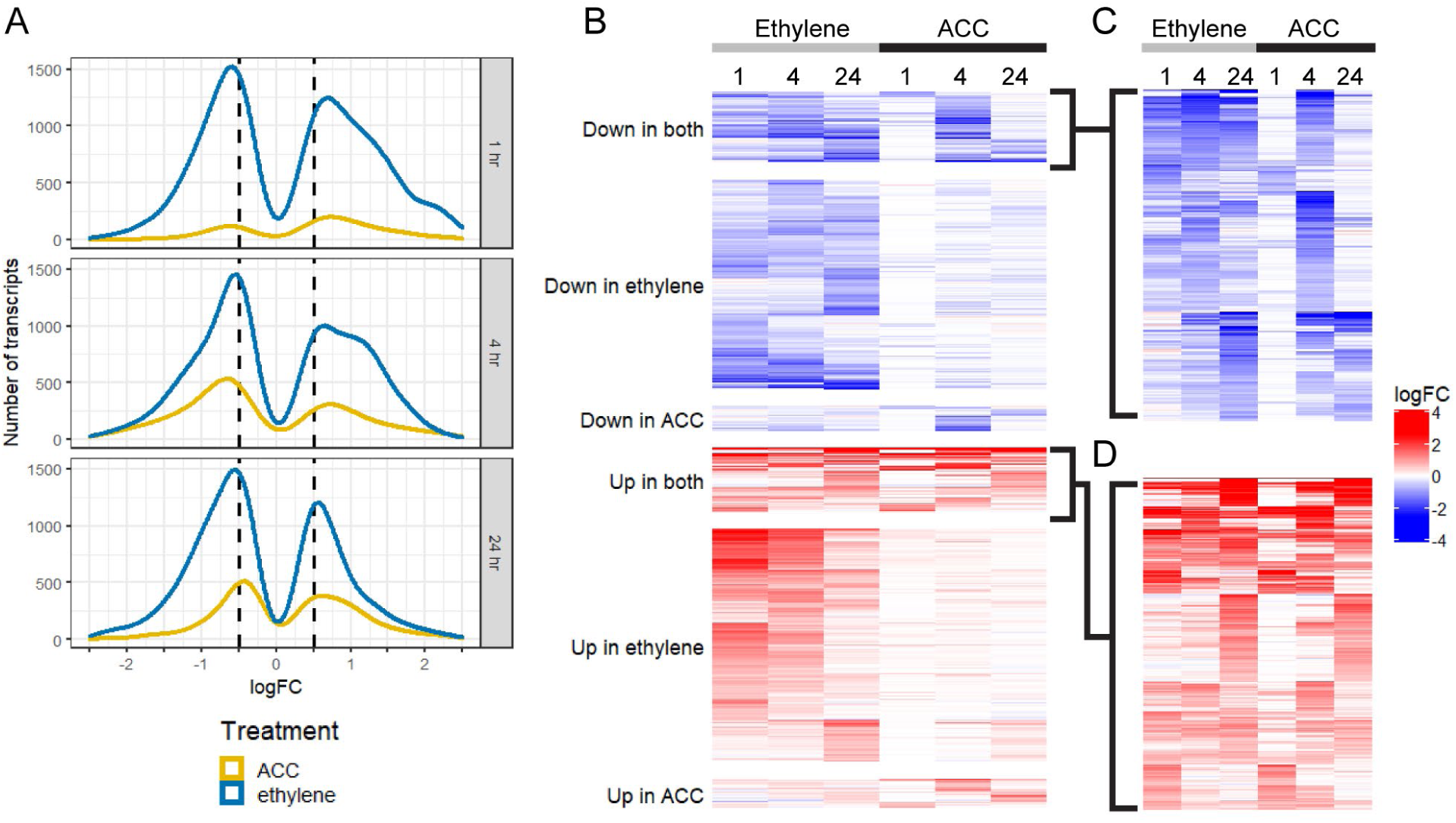
The transcriptional responses to ethylene and ACC treatment revealed more down-regulated genes at 4 and 24 hours and more DE genes by ethylene than ACC. (A) Density plot reporting the number of transcripts across a range of log_2_FC values for each treatment and time point in Col-0 samples. The log_2_FC is relative to Col-0 control (time 0). For each treatment and time point, only transcripts with an adjusted p-value < 0.01 are shown; no log_2_FC cutoff was used. (B) Heatmap showing the log_2_FC values of the 4,522 transcripts that were DE relative to untreated Col-0 in at least one Col-0 sample treated with ethylene or ACC based on |log_2_FC| > 0.5 and adjusted p-value < 0.01. (C & D) Transcripts which are DE in at least one time point for both ethylene and ACC, with a (C) negative or (D) positive log_2_FC.

To identify genes with the most robust changes, we utilized a |log_2_FC| > 0.5 and adjusted p-value < 0.01 to define differential expression (DE) for the following analyses. Using DESeq2, we identified 4,522 transcripts that were DE in response to ethylene and/or ACC in at least one time point in Col-0 (Figure 3B-D). These transcripts and their log_2_FCs (treated Col-0 relative to time-0 Col-0) are listed in Supplemental File 1. More transcripts responded to ethylene treatment and had a greater log_2_FC than to ACC treatment. This log_2_FC cutoff eliminated more genes whose transcripts decreased after treatment, which resulted in a similar number of up- and down- regulated transcripts passing the log_2_FC cutoff, despite there being more down-regulated genes in the larger set of those with a p-value < 0.01 (Figure 3A). Many genes showed similar trends in response to ACC and ethylene treatment: 1,493 and 1,657 genes were down- and up-regulated, respectively, by ethylene only, and 182 and 213 genes were down- and up-regulated, respectively, by ACC only as summarized in Supplemental Table 2. Strikingly, only 16 genes had opposite responses to the two treatments: three were down-regulated in ethylene and up- regulated in ACC, and 13 were up-regulated in ethylene and down-regulated in ACC. This very small number of oppositely regulated genes by these two treatments is consistent with ACC and ethylene leading to common transcriptional responses, which may differ in magnitude of response, but infrequently in the direction of responses.

This analysis also revealed that 457 and 504 transcripts were up- and down- regulated, respectively, by both ACC and ethylene, with heatmaps illustrating the transcript abundance over time in Figure 3C and D. One pattern that became apparent was that for transcripts whose abundance decreased after both treatments, the 1- and 24-hour ACC-treated samples did not respond as strongly as the time-matched ethylene-treated samples, while in the 4-hour ACC treatment, the number of transcripts changed more dramatically than time-matched ethylene treatment. This finding suggested the possibility that the rate of conversion of ACC to ethylene resulted in varied ethylene concentrations across this time course. Indeed, of the 457 transcripts up-regulated by both ACC and ethylene, 258 had a larger magnitude change in response to ACC treatment at the 4-hour time point, but only 70% of them had a larger response to ethylene treatment in the 1-hour time point. Similarly, of the 504 transcripts down-regulated by both ACC and ethylene, 245 had a larger magnitude change in response to ACC treatment at the 4-hour time point, with 87% of them responding more strongly to ethylene treatment at the 1-hour time point. These results suggested that ACC induced maximal transcriptional responses more slowly than ethylene, perhaps due to the time required to convert ACC to ethylene. Intriguingly, the pattern of highest absolute log_2_FC at the 4-hour time point was also apparent in the transcripts that were DE only in response to ACC treatment.

### Transcripts regulated by ethylene and ACC were enriched in functions related to root development

To determine which biological processes were enriched among the transcripts whose abundance changed after ethylene and/or ACC treatment in Col-0, we performed Gene Ontology (GO) analysis using AgriGO (Du et al., 2010; Tian et al., 2017) and reported the results in Supplemental Table 3. Genes whose transcripts had increased abundance in both treatments were enriched in functional annotations including “negative regulation of the ethylene-mediated signaling pathway,” which is consistent with both ethylene and ACC transcriptionally regulating components of the ethylene signaling pathway in a negative feedback loop, and response to auxin stimulus, consistent with crosstalk between ethylene and auxin.

Transcripts whose abundance decreased after ethylene and/or ACC treatment were enriched in annotations pertaining to lignin biosynthesis and cell wall loosening, while transcripts that decreased after ethylene but not ACC treatment were enriched in pathways related to secondary cell wall biogenesis, plant-type cell wall organization, and lignin metabolism. These results are consistent with the requirement for cell wall remodeling during apical root growth, lateral root emergence, and root hair development.

Interestingly, genes whose transcripts decreased in abundance with ethylene treatment only were enriched in annotations of root hair cell differentiation or root hair development, including two myosin genes (*XIB* and *XIK*), *MORPHOGENESIS OF ROOT HAIR 1, 2*, and *6*, *ROOT HAIR DEFECTIVE 4* (*RHD4*), *BRISTLED 1*, *CAN OF WORMS 1* (*COW1*), and *SHAVE 2* and *3*. No annotations were significantly enriched in genes down-regulated by ACC only, likely due to the smaller size of this group.

We further examined the identities of transcripts that were significantly regulated by ACC but not by ethylene, as these transcripts might reveal ethylene-independent, ACC-mediated signaling mechanisms in roots. Among the 213 transcripts that were uniquely up-regulated by ACC were transcripts encoding two mitogen-activated protein (MAP) kinases MAPKKK13 and MEK1, enzymes that remodel the cell wall, such as EXPANSIN A17 (EXPA17) and PECTIN METHYLESTERASE 3 (PME3), and transcripts encoding proteins that function in root hair development, such as RHO OF PLANTS GUANINE NUCLEOTIDE EXCHANGE FACTOR 12 (ROPGEF12) and ROOT HAIR DEFECTIVE 6-LIKE 5 (RSL5). Among the 182 transcripts that were uniquely down-regulated by ACC were *EXPANSIN A4* (*EXPA4*) and *EXPANSIN-LIKE A2* (*EXLA3*). These genes suggest testable hypotheses on mechanism for ACC-regulated root development including cell wall remodeling, MAP kinase signaling, and membrane trafficking.

### Transcriptional responses are accentuated in constitutive-signaling *etr1-7* and reduced in ethylene-insensitive *etr1-3*

We asked how the transcriptional responses to ethylene and ACC varied in the *etr1-3* and *etr1-7* mutants compared to Col-0. Using a cutoff of |log_2_FC| > 0.5 and adjusted p-value < 0.01, we compared each time point for each treatment and genotype to the Col-0 control and found 8,323 transcripts that were DE in at least one of these genotype comparisons, and these transcripts and their log_2_FC values relative to time-0 Col-0 are listed in Supplemental File 1. A heatmap displaying the changes in abundance of these transcripts in all three genotypes and both treatments are shown in Figure 4. All log_2_FC values represented here were relative to Col-0 at time 0 to accurately reflect the changes in baseline levels in *etr1-7*. The first striking observation from this heat map is almost all the DE genes show the same directional change in response to both ACC and ethylene, yet the magnitude of the response is larger in response to ethylene than to ACC for most genes. Additionally, this heatmap hints at the muted response to treatment in *etr1-3* and the enhanced transcript abundance independent of treatment in *etr1-7*.

**Figure 4.**
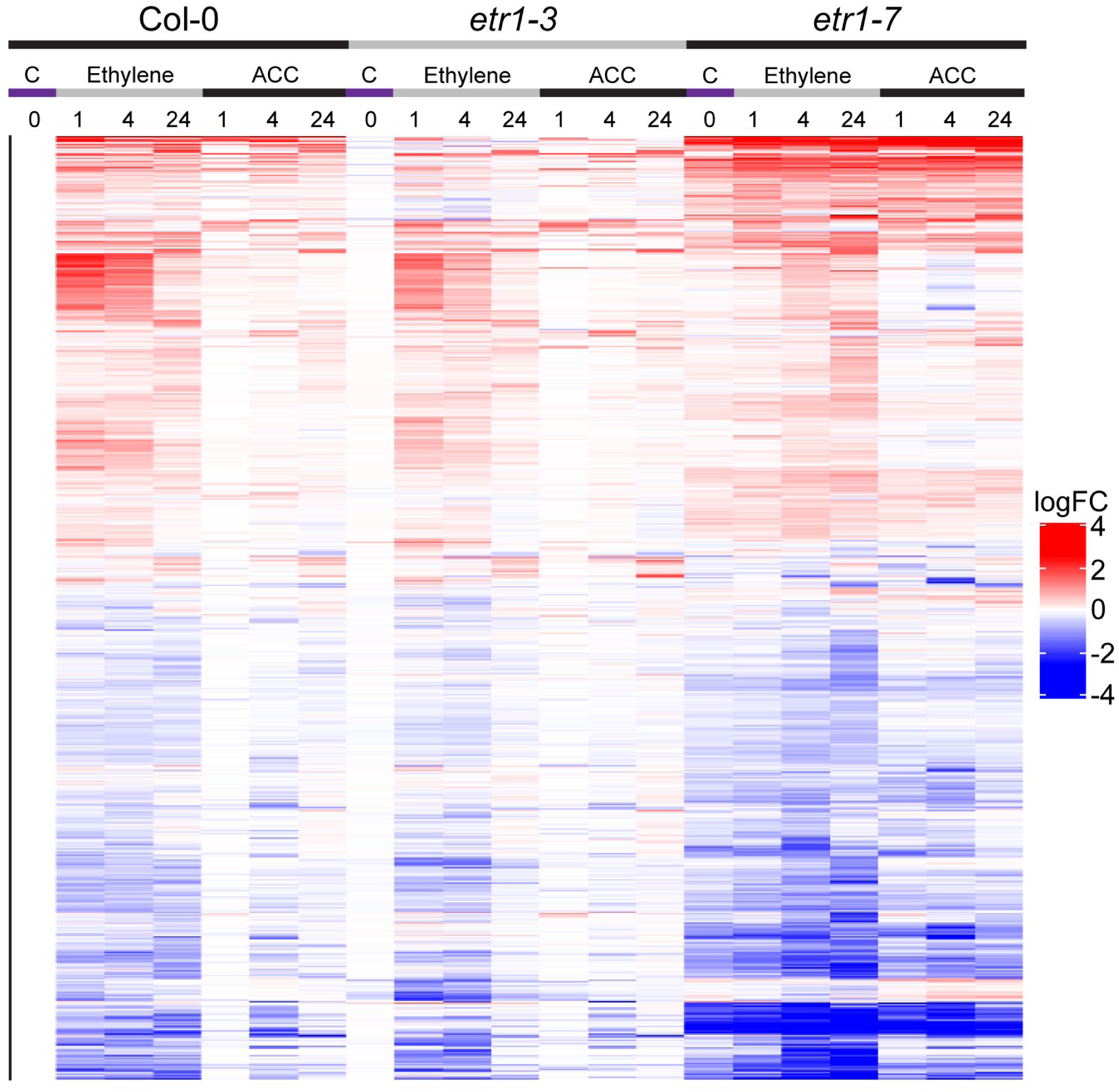
Heatmap of all transcripts that were DE in at least one time point, treatment, or genotype revealed a variety of patterns. This comparison of the transcriptional responses of 8,323 DE transcripts across treatments, time points and genotypes revealed suppressed ethylene signaling in *etr1-3* and enhanced ethylene signaling in *etr1-7*, relative to Col-0 untreated. The heat map includes all 8,323 transcripts that are DE in at least one of the represented groups in comparison with Col-0 control with |log_2_FC| > 0.5 and adjusted p-value < 0.01. log_2_FC for all columns is relative to untreated Col-0 samples.

The heatmap in Figure 4 suggested that the log_2_FC filter eliminated transcripts that responded to ACC but had smaller magnitude changes in response to ACC than ethylene. As illustrated in the density plots, many transcripts had a |log_2_FC| between 0.25 and 0.5, so we examined transcriptional responses using a less stringent cutoff as detailed in the methods. Regardless of cutoff, the number of transcripts that only responded to ethylene was greater than those only responding to ACC, as shown in Supplemental Table 3, where the |log_2_FC| > 0.25. We therefore used the more stringent cut off in our experiments.

We compared baseline levels of expression in time-0 samples between genotypes (Figure 5A), identifying 2,562 genes that were DE (p-value < 0.01) in at least one of these comparisons (these transcripts and their log_2_FC values relative to other genotypes are listed in Supplemental File 1). Consistent with our expectations, *etr1-3* and Col-0 were nearly identical in the absence of elevated ethylene; only 49 out of 2,562 genes were found to be DE between them. However, when we compared *etr1-7* with either Col-0 or *etr1-3*, we found a dramatic difference: 1,941 genes were DE between *etr1-7* and Col-0, and 2,078 genes were DE between *etr1-7* and *etr1-3*. A large number (1,458) of the genes were DE in both comparisons with *etr1-7,* including genes enriched in the “response to ethylene” annotation (Figure 5B).

**Figure 5.**
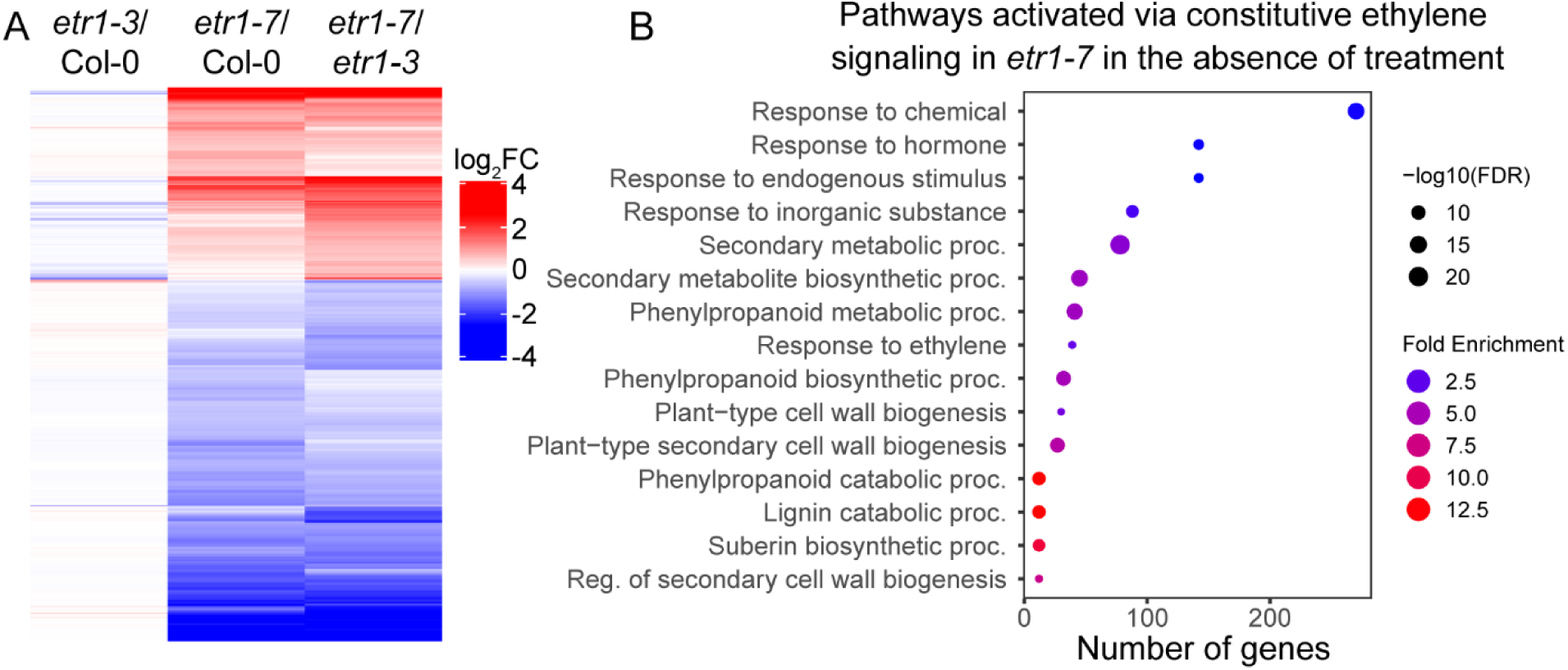
Baseline transcript abundance differences between Col-0 and the *etr1* mutants were consistent with constitutive ethylene signaling in *etr1-7*. (A) All transcripts which are DE in at least one comparison with |log_2_FC| > 0.5 and adjusted p-value < 0.01. log_2_FC is relative to the second genotype in each comparison. (B) Enriched biological processes among the 1,458 DE genes in the overlap between the *etr1-7* vs. Col-0 comparisons and the *etr1-7* vs. *etr1-3* comparisons as determined by ShinyGO 0.80 (Ge et al., 2020).

One very interesting group of transcripts were the 1,941 transcripts that were DE in *etr1- 7* as compared to Col-0, in the absence of hormone treatment. The large number and magnitude change of these genes is consistent with the *etr1-7* mutation being sufficient to lead to large changes in abundance of transcripts in the presence of endogenous levels of ethylene due to enhanced ethylene signaling. Under these conditions, the *etr1-7* mutant has increased root hair formation relative to Col-0. Consistent with this group of genes having an important function in ethylene response, this group was enriched in annotations pertaining to the ethylene signaling pathway. Similar annotations were also found in the group of transcripts that responded to ethylene and/or ACC treatment in Col-0, including “response to ethylene” (Supplemental Table 3). These results are consistent with *etr1-7* having nearly constitutive ethylene signaling and therefore a different transcriptional landscape than the other genotypes in the absence of exogenous ethylene or ACC and with transcript abundance changes that overlap with Col-0 after treatment with ACC or ethylene.

### A subset of ACC- and ethylene-regulated transcript responses are dependent on ETR1

We developed a rigorous set of criteria to determine which ethylene- and/or ACC-regulated transcripts were dependent on ETR1 for transcriptional response. For this analysis, we included transcripts that were DE in response to ACC or ethylene designated as a |log_2_FC| > 0.5 and a p-value < 0.01 and defined transcripts as ETR1-dependent based on the following criteria: 1) the transcript was DE with ethylene and/or ACC treatment in at least one time point in Col-0; 2) the transcript showed no significant change in any *etr1-3* samples compared to Col-0 time 0; and 3) the transcript was DE in *etr1-7* relative to Col-0 in the baseline and showed similar abundance in ACC- and ethylene-treated *etr1-7* samples. Since *etr1-7* has constitutive ethylene signaling, it should have had transcriptional responses that differed from Col-0 in the absence of either ACC or ethylene treatment and were unchanged by ACC or ethylene treatment. Therefore, we compared *etr1-7* treated samples to *etr1-7* control samples, and only those that did not change with treatment in *etr1-7* qualified as ETR1-dependent.

With this strict criteria, 553 transcripts were identified as ETR1-dependent, which are shown in the heat map in Figure 6 (and Supplemental File 1) with their log_2_FC values relative to time-0 Col-0). A subset of 292 transcripts met these criteria in ethylene treatment only, another 191 in ACC only, and 70 genes were classified as ETR1-dependent in both treatments. The patterns in this heat map are striking leading to four major conclusions. First, it is clear that the *etr1-7* mutant leads to profound changes in transcript abundance (relative to time-0 Col-0) even without treatment, which is evident in the time-0 *etr1-7* sample (labeled C), which is normalized to the Col-0 time-0 sample) and that there are profound increases that are independent of treatment. Second, there is a muted response in *etr1-3* that is evident for treatment with either ethylene or ACC. Third, the response of these ETR1-dependent transcripts to ethylene is generally greater than to ACC, which may be tied to potency of doses of these two molecules, although ACC induced a larger effect at the 4-hour time point in Col-0, which is highlighted in a green box on the heatmap. Fourth, the larger number of DE genes in response to ethylene than ACC may be tied to a smaller magnitude of ACC response that was below the log_2_FC cutoffs, suggesting that these transcripts are regulated by ETR1-dependent in both treatments. We also examined the transcripts filtered to have a |log_2_FC| > 0.25, which yielded many more transcripts in each group, but did not change the overall ratios of transcripts in those groups.

**Figure 6.**
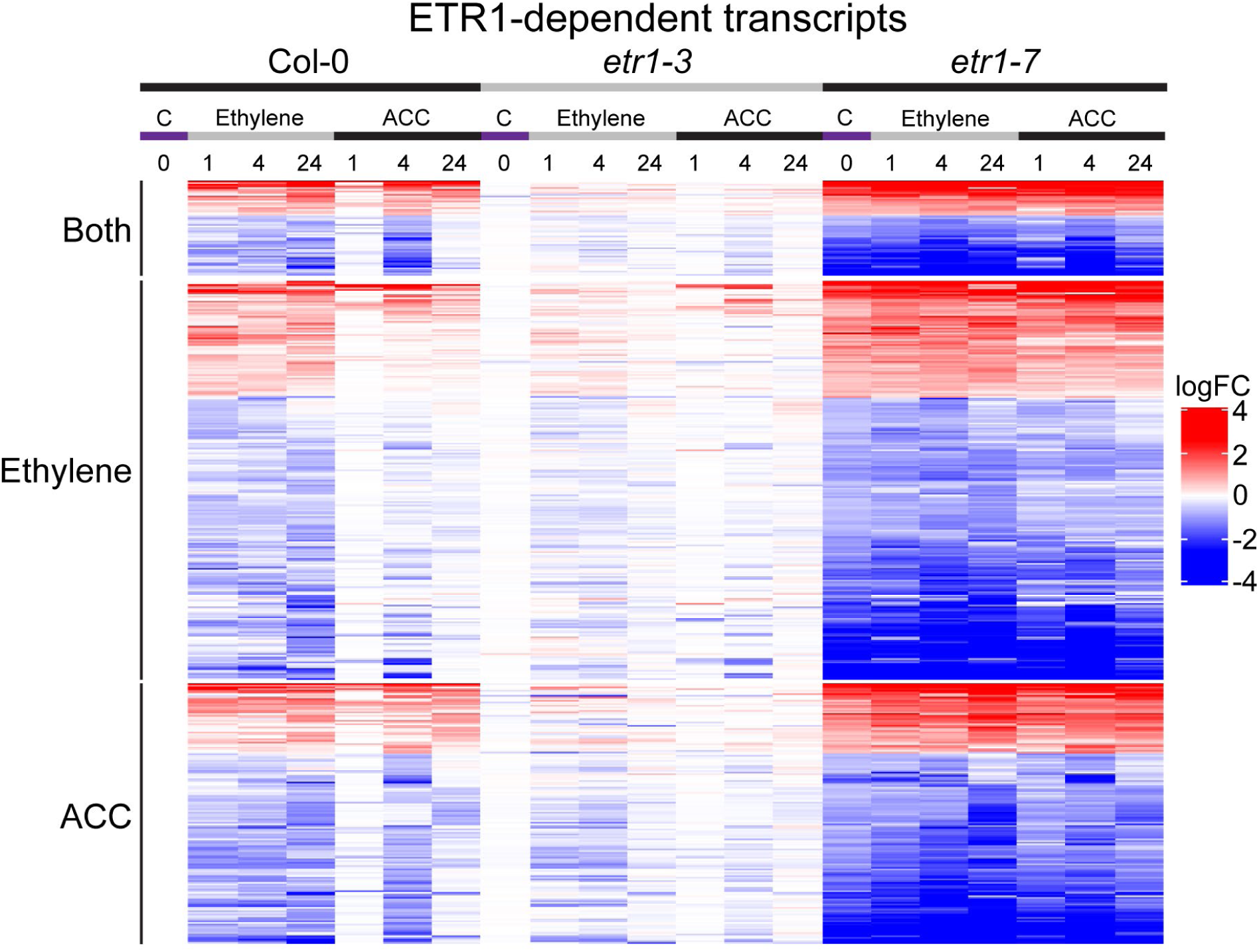
ETR1-dependent transcript heatmap revealed genotype and treatment dependent changes in transcript abundance. This comparison of transcript abundance across treatments and genotypes revealed a set of transcripts that responded to ethylene and/or ACC treatment in Col-0 but not in the mutants. Transcripts which follow a pattern of response that suggests they are ETR1-dependent were selected. These transcripts respond to ACC and/or ethylene treatment in Col-0, do not respond in *etr1-3*, and are constitutively up or down in *etr1-7*, based on |log_2_FC| > 0.5 and adjusted p-value < 0.01. The log_2_FC reported here was determined relative to untreated Col-0. Genes are split into three groups based on whether they met our ETR1-dependent criteria in ethylene only (292 transcripts), ACC only (191 transcripts), or both treatments (70 transcripts).

Out of the 553 genes which qualified as ETR1-dependent in one or both treatments, 31 overlapped with a set of core ethylene-responsive genes identified by a comparison of three transcriptome studies of ethylene- or ACC-treated roots (Harkey et al., 2019). These genes included *ACC OXIDASE 2* (*ACO2*), *ETHYLENE RESPONSE SENSOR* (*ERS)1* & *2*, and *ETHYLENE AND SALT INDUCIBLE 3* (*ESE3*). This finding suggested there were ETR1- dependent transcriptional responses to ACC and ethylene in this core group. Other ethylene-related genes that did not overlap with this group but were identified as ETR1-dependent here included *ACO3*, several *ERFs*, and *PINOID*, a kinase that is a positive regulator of auxin transport (Sukumar et al., 2009) and a negative regulator of root hairs (Lee and Cho, 2008). The 553 ETR1-dependent transcripts were enriched in GO annotations including lipid transport, response to ABA, and regulation of transcription (Supplemental Table 4).

We tested an additional set of less stringent criteria, where response to treatment was evaluated by comparing treated samples to the control samples within each genotype, but only required presence or absence of treatment response regardless of the control sample abundance. In this case, ETR1 dependence was defined by a response in Col-0 but not in the mutants, and ETR1 independence was defined as a response in all three genotypes. This analysis allowed for a larger set of transcripts to be included in both groups (Supplemental Figure 5), but ultimately the changes in abundance of the additional transcripts were not as strong as those seen with the more stringent criteria, so we focused our analysis on the transcripts that met the more stringent criteria.

### Other transcripts appear to be regulated by ACC and/or ethylene in an ETR1-independent or more complex manner

We defined ETR1-independent responses for each treatment as having a response to treatment in all genotypes and no altered response in the mutants at baseline (compared to time-0 Col-0). We compared the *etr1-7* treated samples to the *etr1-7* control samples to determine whether a gene responded to treatment in this genotype relative to baseline transcript levels, but in this case a response to treatment was required to qualify as ETR1-independent.

The search for ETR1-independent genes resulted in a slightly smaller set of genes: 481 genes qualified as ETR1-independent in ethylene only, 95 in ACC only, and 117 qualified in both treatments (listed in Supplemental File 1 with their log_2_FC values relative to time-0 Col-0). These ETR1-independent transcripts are shown in the heatmap in Figure 7, where the log_2_FC relative to time-0 Col-0 was reported. What is very striking in this figure is the consistent pattern of the genes that are DE in response to both ethylene and ACC show no difference in response in all 3 genotypes. This trend is also true in the ethylene-only DE group, but the ACC response is muted. The ACC only group had a very sporadic pattern with multiple different responses. Another interesting observation was found among the ETR1-independent transcripts regulated by both ACC and ethylene. Transcripts that were up-regulated by both treatments were rapidly induced at the 1-hour time point, while transcripts that were down-regulated were more slowly suppressed at the 4-hour time point. GO annotations enriched in this group of genes included several stress-response annotations, as well as “root hair cell differentiation” and “unidimensional cell growth” (Supplemental Table 4).

**Figure 7.**
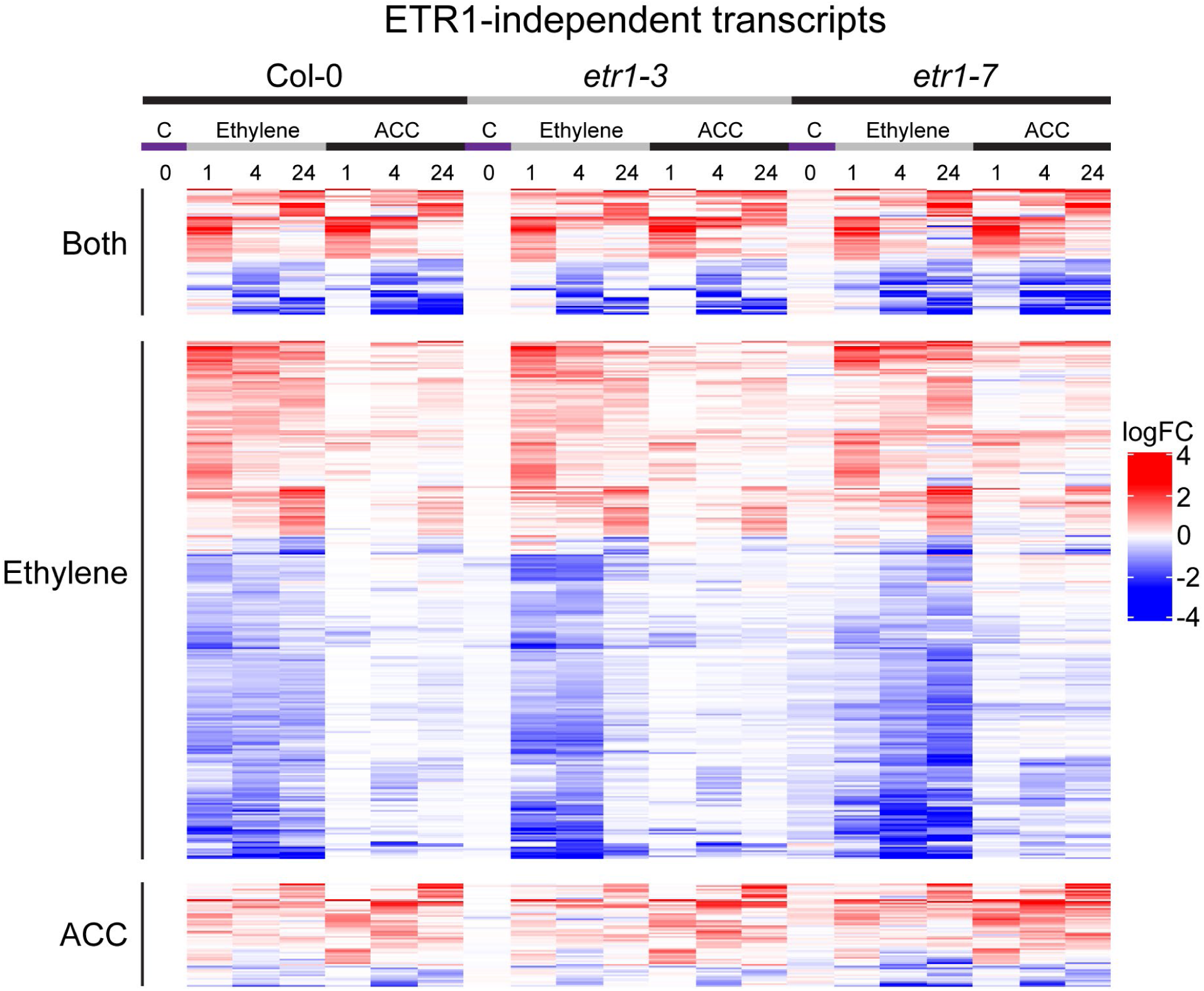
ETR1-independent transcript heatmap revealed genotype and treatment independent changes in transcript abundance. This comparison of transcript abundance across treatments and genotypes revealed a set of genes that responded to ethylene and/or ACC treatment in all genotypes. Genes which respond to ethylene or ACC in all three genotypes were considered receptor-independent (since transcript abundance should have been constant in *etr1-3* and *etr1-7)*. Significant differences were defined as having |log_2_FC| > 0.5 and adjusted p-value < 0.01. Transcripts are divided based on which treatment qualifies as ETR1-independent. log_2_FC is reported relative to untreated Col-0. Genes are split into three groups based on whether they met our ETR1-independent criteria in ethylene only (488 transcripts), ACC only (95 transcripts), or both treatments (117 transcripts).

After grouping of genes into ETR1-dependent and -independent responses, there were an additional 7,077 genes that were DE in response to ethylene- or ACC-treatment in at least one sample but did not meet the criteria to fit in either group (listed in Supplemental File 1 with their log_2_FC values relative to time-0 Col-0). These genes did not meet the strict criteria defined for ETR1-dependent or -independent genes above, showing patterns of response inconsistent with either definition (Supplemental Figure 6).

We also compared our ACC regulated genes to a recent dataset where *ein2-5* seedlings were treated with 10 µM ACC for 40 hours to identify ACC-regulated transcripts (Mou et al., 2025). This study identified 211 DE genes that were down-regulated and 147 that were up- regulated by ACC. We asked whether these DE genes were also regulated by ACC in our root- specific dataset, finding that, indeed, a majority of the up-regulated genes were (108 out of 147), although only 17 out of 212 down-regulated genes were ACC regulated in our dataset, but over 100 of these were ethylene-regulated. We also asked if their DE genes were found in our ETR- independent dataset, as the phenotypes of *etr1-3* and *ein2-5* mutants are similar. None of their up-regulated genes were DE in response to ACC treatment in the ETR1-independent dataset, while 32 of their up-regulated genes (22%) were ACC regulated in our ETR1-independent dataset. The limited overlap between these two ACC-treatment datasets was unexpected, but these differences could be due to seedling age, genotype, growth conditions, time of treatment, and isolation of RNA from whole seedling versus roots.

### Several ACOs are transcriptionally regulated by ACC and ethylene with some dependent on the ETR1 receptor

An important action of both ACC and ethylene is to positively regulate the synthesis of more ethylene. In particular, the ACC oxidase (ACO) enzyme catalyzes the oxidation of ACC to ethylene as the final step of ethylene biosynthesis, which may use either endogenously synthesized and exogenously applied ACC as a substrate. Included in the ETR1-dependent dataset were transcripts encoding several of the five ACOs (*ACO1* through *ACO5*). The transcript levels of *ACO1*, *ACO2*, and *ACO3* in each genotype and treatment are shown in Figure 8, revealing that transcripts encoding ACO enzymes are positively regulated by ACC and ethylene levels, but with a stronger response to ACC. This ACC-driven increase in ACO enzyme synthesis has the potential to increase conversion of exogenous ACC to ethylene. The abundance of transcripts from these genes varied in time-0 Col-0 roots with higher transcript abundance of *ACO2* and *ACO5* (313 and 315 TPM, respectively), while *ACO1*, *ACO3*, and *ACO4* had TPM values between 6 and 39. We found that both ethylene and ACC treatment of Col-0 significantly increased the transcript abundance of four *ACO*s (*ACO1*, *ACO2*, *ACO3*, and *ACO5*) in at least one time point after treatment. For these genes, there was a greater magnitude increase in transcript abundance in response to ACC than to ethylene treatment in at least one time point for each transcript. This difference in response is striking as most of the up-regulated DE transcripts in our dataset had larger magnitude changes in response to ethylene than to ACC treatment. This finding suggests that exogenous ACC may be a stronger signal to promote transcript accumulation of *ACO*s than ethylene gas itself.

**Figure 8.**
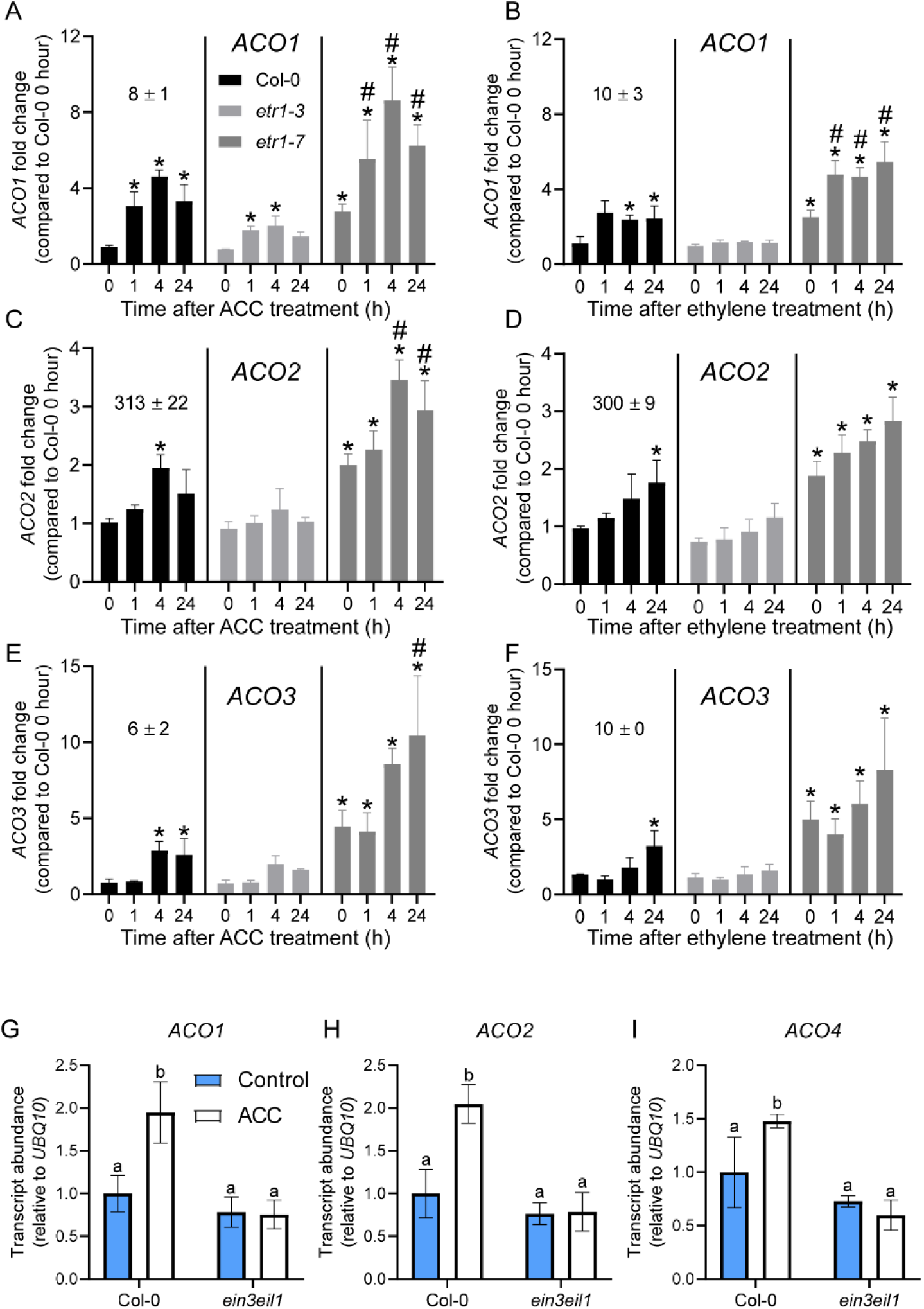
Transcript abundance of ETR1-dependent and/or EIN3/EIL1-regulated *ACO*s. (A-F) Normalized transcript abundance was calculated for *ACO1, ACO2*, and *ACO3* by taking average TPM for each time point and reporting it relative to the average TPM of untreated Col-0. Asterisks indicate statistical significance (p-value < 0.01) compared to Col-0 time 0. Pound symbols indicate statistical significance (p-value < 0.01) compared to *etr1-7* time 0 and are displayed only for *etr1-7*. Error bars represent SD. (G-I) Transcript abundance in Col-0 and *ein3eil1* samples treated with and without 0.75 µM ACC for 4 hours was quantified using RT-qPCR. The average of three biological replicates are reported with error bars representing SD. Statistical significance was defined as p-value < 0.05 using Tukey’s multiple comparisons test.

According to our strict criteria for ETR1 dependence, detailed above, *ACO2* and *ACO3* were ETR1-dependent in response to ethylene treatment, with uniform reductions in transcript abundance in *etr1-3* and increases in *etr1-7* compared to Col-0 (Figure 8D, F). While *ACO2* and *ACO3* passed the criteria for ETR1 dependence in Col-0 and *etr1-3* in response to ethylene treatment, ACC induced significant increases in transcript abundance at one time point in the *etr1-7* mutant relative to the *etr1-7* time-0 control and therefore these *ACOs* did not pass the criteria for ETR1 dependence in the presence of ACC. *ACO1*, *ACO4,* and *ACO5* did not meet our criteria for ETR1 dependence in the presence of either ACC or ethylene (Figure 8A, B; Supplemental Figure 7). *ACO5* appeared to be ETR1-dependent based on p-value, but it failed the log_2_FC cutoff for having a smaller magnitude change than required for our stringent requirement. It is clear from the graphs that reduced or constitutive ethylene signaling in the *etr1-3 and etr1-7* mutants, respectively, altered the transcriptional response to ACC and/or ethylene for each *ACO*, consistent with the importance of the ETR1 receptor in regulating *ACO* transcript abundance, even for *ACO*s that do not meet our stringent ETR1-dependent criteria.

### ACC-induced transcript accumulation of ACO1, ACO2, and ACO4 is mediated by EIN3/EIL1

To extend our understanding of the regulation of *ACO*s, we asked if their changes in transcript abundance in roots were regulated by both ETR1- and EIN3-mediated GRNs. We quantified the transcript abundance of the *ACO*s in roots of Col-0 and *ein3eil1* using RT-qPCR in the presence and absence of 0.75 µM ACC. We treated these samples with ACC for 4 hours, as this was the time point that yielded the largest increases in *ACO* transcript abundance (relative to time-0 Col-0) in our RNA-Seq data. For *ACO1*, *ACO2*, and *ACO4*, we found significant increases in transcript levels with ACC treatment in Col-0, including 2-fold increases in *ACO1* and *ACO2* and a 1.5-fold increase in *ACO4* (Figure 8G-I). These ACC-induced increases were lost in the *ein3eil1* mutant, consistent with these transcripts being ACC-regulated through the canonical ethylene signaling pathway (Figure 8G-I). Consistent with our RT-qPCR results, the promoters of *ACO2 and ACO4* were found to be directly bound by EIN3 in a DAP-Seq (O’Malley et al., 2016) and/or ChIP-Seq dataset (Chang et al., 2013) as summarized in Harkey et al. 2019, but the absence of *ACO1* as an EIN3 target is surprising and is consistent with an EIN3- regulated TF binding to and controlling *ACO1* transcription, rather than being a direct EIN3 target.

In contrast, we did not detect significant increases in *ACO3* and *ACO5* transcripts in response to ACC treatment in Col-0 in our RT-qPCR (Supplemental Figure 8). *ACO3* transcript levels in *ein3eil1* did not differ from those in Col-0 in the presence or absence of ACC, while *ACO5* transcript levels in Col-0 ACC-treated samples were significantly higher than those in *ein3eil1* in the presence and absence of ACC (Supplemental Figure 8). For both *ACOs*, these findings were consistent with localized changes in abundance in specific tissues that cannot be consistently detected by RT-qPCR using whole roots.

### ACO transcription increases in distinct root cell types in response to ACC treatment

To determine which of the *ACO*s might control ACC- and ethylene-induced changes in root development, we queried the *ACO*s in a root cell-type-specific dataset (Brady et al., 2007) to identify the tissues where these transcripts were most highly enriched. We searched for transcripts that were localized in the appropriate tissues to mediate ACC- and ethylene-induced root responses, including stimulation of root hair initiation, inhibition of lateral root emergence and inhibition of root elongation at the root tip. *ACO5* was identified as a candidate for the control of root hair growth because, like *ETR1*, its transcripts were highly enriched in cells expressing root hair markers, while the other *ACOs* had low transcript abundance in this cell type (Figure 9A).

**Figure 9.**
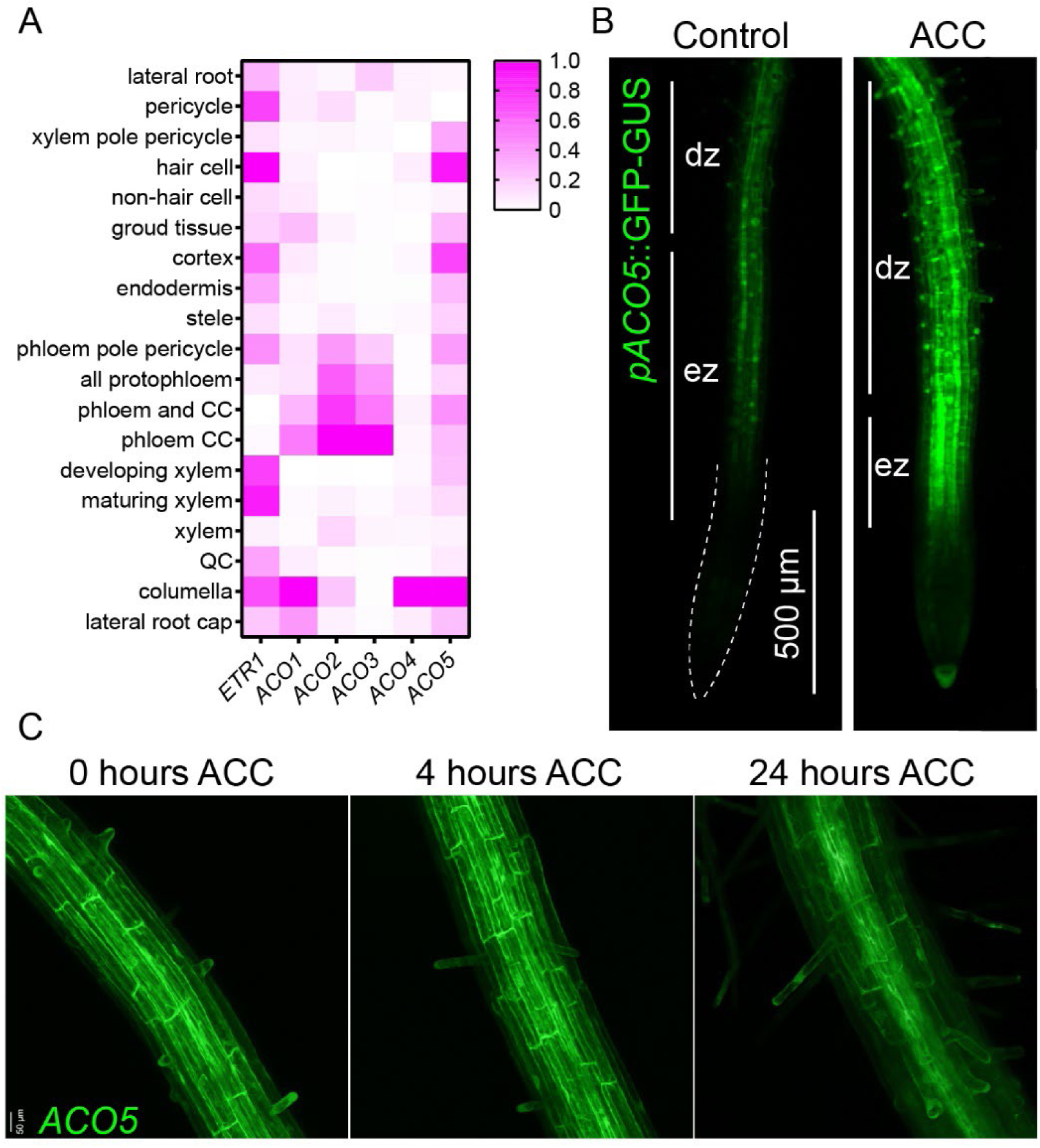
*pACO5*::GFP-GUS is expressed in roots including root hairs. (A) Publicly available root cell type data reported as a heatmap (Brady et al., 2007). Data has been scaled and normalized to the tissue with the highest expression for each transcript. (B) Roots of five-day-old *pACO5*::GFP-GUS seedlings treated with and without 0.75 µM ACC for 4 hours with images captured on a fluorescent stereomicroscope. DZ, differentiation zone. EZ, elongation zone. Scale bar is 500 µm. (C) Roots of 5-day-old *pACO5*::GFP-GUS seedlings treated with 0.75 µM ACC for the indicated time were imaged by laser scanning confocal microscopy. Scale bar = 50 µm.

To provide additional insight into the effect of ACC treatment on tissue-specific localization and abundance of *ACO* transcripts, we examined a fluorescent reporter driven by each of the native *ACO* promoters (Houben et al., 2024). To validate the root hair cell specific *ACO5* accumulation in the microarray described above, we examined fluorescence of a *pACO5*::GFP-GUS reporter in roots. Comparison of root tip with and without a 4 hour ACC treatment reveals low levels of GFP signal at the root tip with higher levels in the elongation and differentiation zones. The GFP signal is evident in hair cells and root hairs, which are increased in number after ACC treatment (Figure 9B). Laser scanning confocal microscope (LSCM) images of root tips of 5-day-old seedlings revealed brighter fluorescence in hair cells than non- hair cells, consistent with cell-type-specific expression patterns (Figure 9B; Figure 10). Treatment with ACC for 4 and 24 hours (Figure 9C) or 5 days (Supplemental Figure 9) led to increased number and elongation of root hairs, further illustrating the presence of *pACO5* driving GFP fluorescence in these cells (Figure 9B). These results indicate that *ACO5* is expressed in the appropriate position for its protein product to synthesize ethylene to drive root hair elongation.

**Figure 10.**
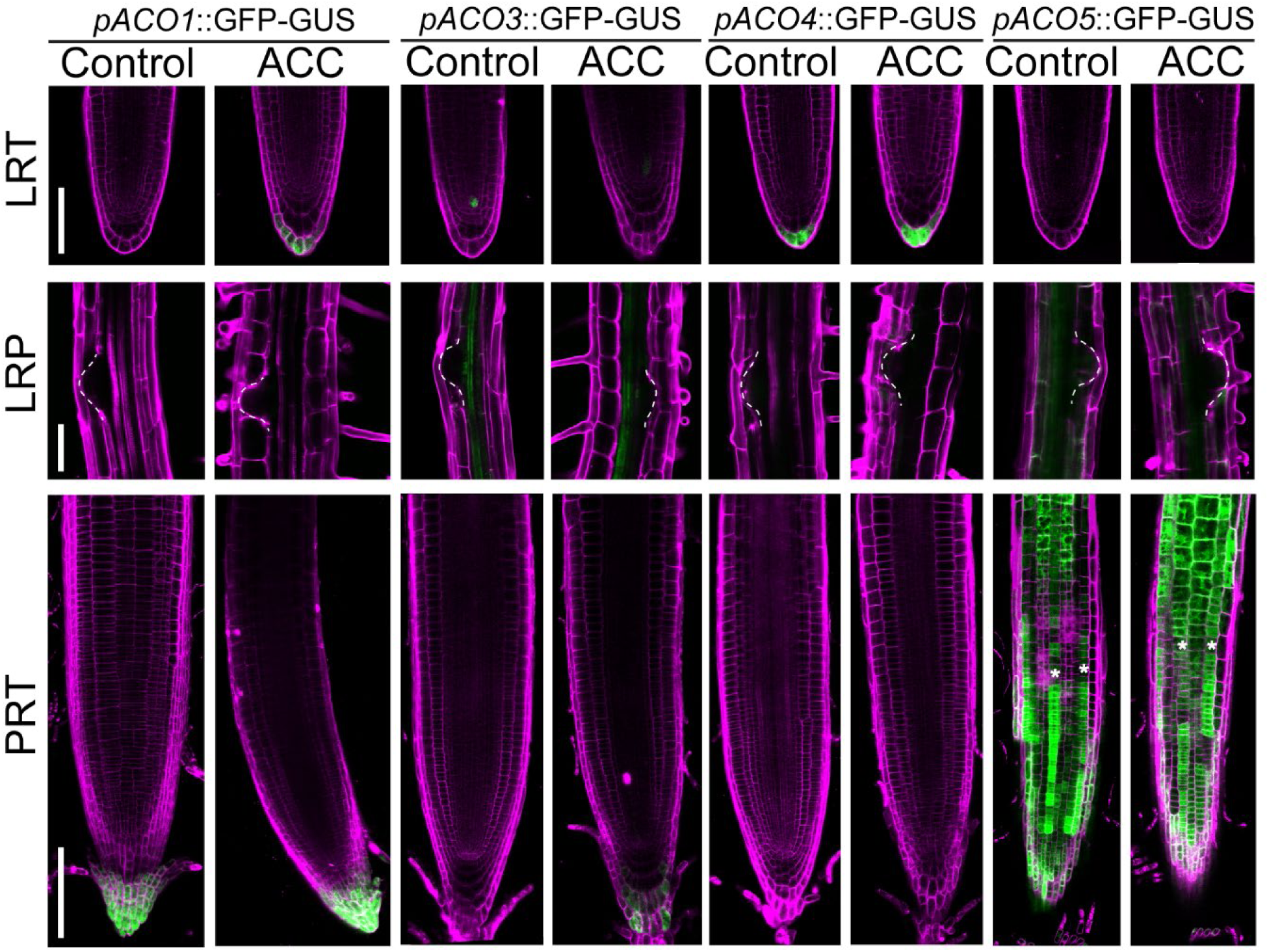
*ACO* transcripts are synthesized in distinct root tissues in response to ACC treatment. 10-day-old *pAC*O::GFP-GUS seedlings treated with and without 0.75 µM ACC for five days. Abbreviations for root regions are primary root tip (PRT), lateral root primordium (LRP), and lateral root tip (LRT). Scale bar is 150 µm for PRT and LRT images and 100 µm for LRP images.

We found shared and distinct ACO expression patterns in roots of 10-day-old seedlings in the presence and absence of 0.75 µM ACC after five days of treatment (Figure 10). The *pACO1*::GFP-GUS reporter had a fluorescent signal in the lateral root cap and the columella of primary and lateral root tips, where this signal was increased in response to ACC treatment. The *pACO3*::GFP-GUS reporter exhibited a fluorescent signal in the primary root columella and the inner tissues of primary and lateral roots, where this signal was increased in response to ACC treatment. Interestingly, the *pACO4*::GFP-GUS reporter had a fluorescent signal in the columella of elongated lateral roots that was increased after ACC treatment, but this signal was not detected in the columella of primary root tips in the presence or absence of ACC. We identified weak ACC-induced changes for some of the reporters in the root tissues of interest, although there appeared to be moderate basal levels of expression consistent with the cell type data (Figure 10). The fluorescence of the *pACO2*::GFP-GUS reporter was low and did not show induction by ACC in root tissues of interest (Supplemental Figure 10). These diverse expression patterns of the *ACO* reporters suggest that the conversion of ACC to ethylene may be controlled at the tissue level locally elevate ethylene signaling.

### Identification of EIN3-regulated transcripts predicted to encode TFs within the ETR1- dependent GRN

One goal of this study was to gain additional insight into the transcriptional regulators downstream of ETR1 whose abundance changed in response to ACC and ethylene treatment. Consistent with a global remodeling of the transcriptome, a GO analysis revealed that the ETR1- dependent transcripts were significantly enriched in genes that function in the regulation of transcription (Supplemental Table 4). We identified 60 ETR1-dependent transcripts predicted to encode TFs using the AGRIS database (Palaniswamy et al., 2006). The expression patterns of these TF-encoding transcripts were calculated and the log_2_FC relative to time-0 Col-0 is shown in Figure 11A. Note the larger number of down-regulated than up-regulated transcripts and the distinct temporal responses, with some transcripts changing rapidly after ACC and/or ethylene treatment and other transcripts responding more slowly to these treatments.

**Figure 11.**
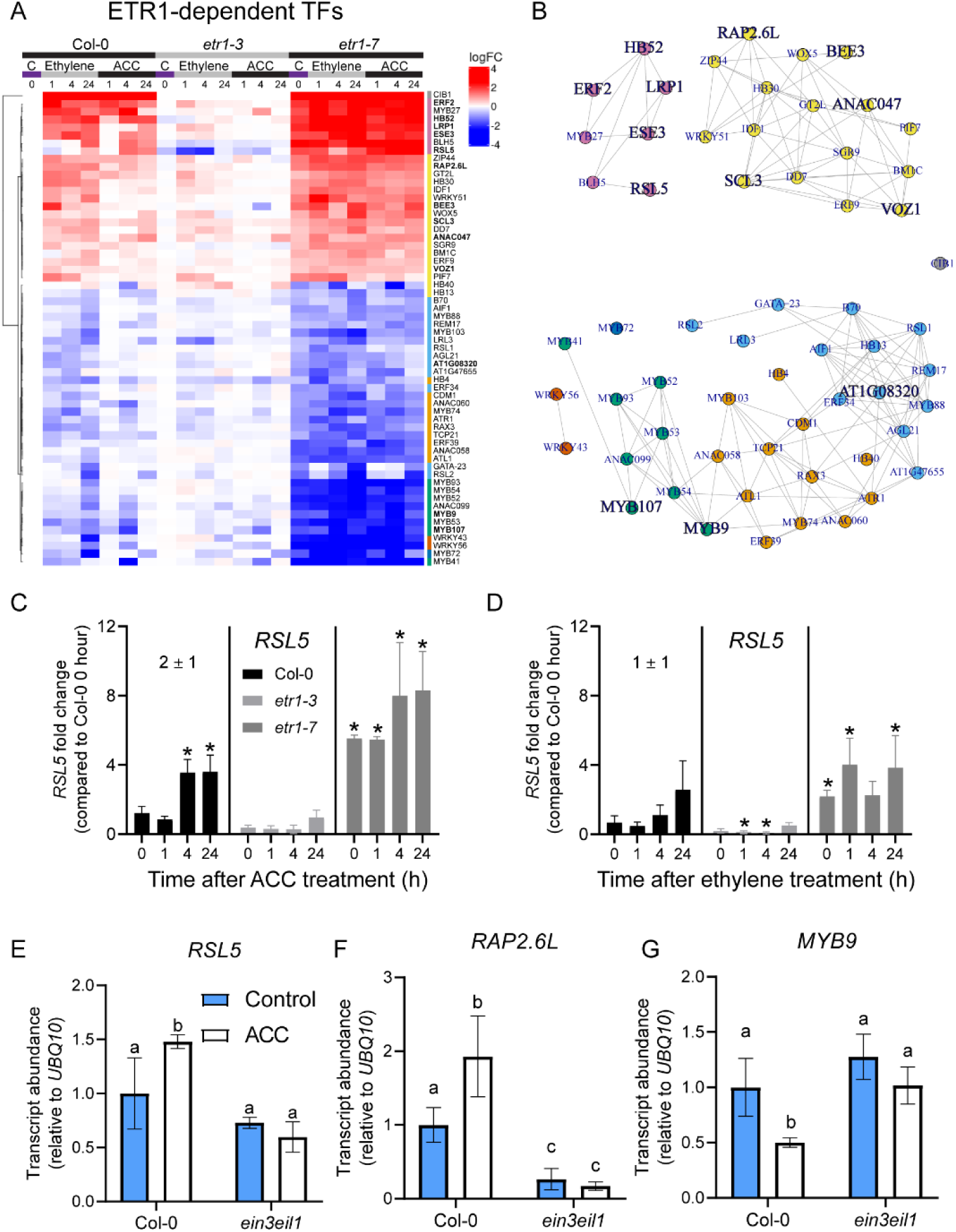
ETR1-dependent transcripts annotated as transcription factors (TFs) exhibit diverse expression patterns. (A) Heatmap of transcript abundance of the 60 ETR1-dependent transcripts that are predicted to encode TFs (reported as log2FC relative to Col-0 time 0). Colors next to gene names correspond to colors in the PaLD network. (B) PaLD network of the ETR1- dependent TFs, with EIN3 direct targets highlighted on the network in large, bold font. The node for CIB1 was positioned more closely to the main network to conserve space. (C, D) RSL5 normalized transcript abundance was calculated by taking average TPM for each time point and reporting it relative to the average TPM of untreated Col-0. (E-G) Transcript abundance in the roots of Col-0 and ein3eil1 seedlings treated with and without 0.75 µM ACC for four hours. Error bars represent SD. Statistical significance was defined as P < 0.05 according to Tukey’s multiple comparisons test.

To better visualize groups of TFs that have similar expression patterns in response to treatments and in the *etr1* mutants, we employed a PaLD analysis (Berenhaut et al., 2022; Khoury et al., 2024a) using the log_2_FC values relative to Col-0 time 0 for each transcript to generate a network of the 60 ETR1-dependent TFs (Figure 11B). This analysis results in a network with each node representing one of the 60 TF-encoding transcripts and the edges connecting transcripts with the most similar responses to ACC and ethylene over time in the different genotypes.

To sort these transcripts into distinct groups with the most similar responses, we used the Louvain method, which identifies communities in large networks (Blondel et al., 2008). The network yielded a structure consisting of six groups containing two or more transcripts. Transcripts belonging to the same group as indicated by their matching colors are those with the most similar transcript abundance patterns across the time course of ACC and ethylene treatment in Col-0, *etr1-3*, and *etr1-7*. As EIN3 is a transcription factor that is downstream of ETR1 in the canonical ethylene signaling pathway (Dolgikh et al., 2019), we used publicly available data on the genes to which EIN3 binds revealed by DAP-Seq (O’Malley et al., 2016) or ChIP-Seq (Chang et al., 2013). We highlighted on the PaLD network in large, bold font the direct targets of EIN3. Consistent with EIN3 functioning to primarily increase transcript abundance, most of the targets were found in the up-regulated groups of genes (Figure 11B). We also calculated the percent of genes in each Louvain group that are direct EIN3 targets. The pink and yellow groups making up the up-regulated transcripts consisted of 71% and 31% EIN3 targets, respectively. These numbers are substantially higher than the 4% of the whole genome containing EIN3 targets. In contrast, the green, blue, orange, and red-orange groups consisting of down-regulated transcripts included 25%, 8%, 0%, and 0% EIN3 targets, respectively. The up-regulatedEIN3 targets had diverse expression patterns, with some being rapidly induced (e.g., *ERF2*) or repressed (e.g. AT1G08320) by ACC/ethylene treatment, and others being more slowly up- (e.g., *RSL5*) or down-regulated (e.g. *MYB9* and *MYB107*).

To validate whether both up-regulated and down-regulated ETR1-dependent transcripts that are EIN3 targets by ChIP-Seq and DAP-Seq showed altered transcript abundance in the *ein3eil1* mutant, we quantified the transcript abundance of several genes in roots of five-day-old Col-0 and *ein3eil1* seedlings in the presence and absence of ACC after 4 hours. We were interested in *ROOT HAIR DEFECTIVE 6-LIKE 5* (*RSL5*), as this TF was previously demonstrated to be involved in root hair development (Pires et al., 2013). Intriguingly, in our RNA-Seq dataset, *RSL5* transcript levels were significantly up-regulated by ACC at the 4- and 24-hour time points but were not significantly regulated by ethylene (Figure 11C, D). Consistent with these data, we found a significant increase in *RSL5* transcripts in the presence of ACC and demonstrated the dependence of this change on EIN3, as this induction was lost in the *ein3eil1* mutant (Figure 11E). This result suggested that although *RSL5* was more strongly regulated by ACC, this regulation depended upon the canonical ethylene signaling pathway. We also observed a significant increase in *RAP2.6L* transcript levels and a significant decrease in *MYB9* transcript levels in the presence of ACC, and both changes were lost in the *ein3eil1* mutant (Figure 11F, G). We also examined transcript abundance of an additional gene (*LRP1*) whose promoter was targeted by EIN3, and three genes (*ANAC058*, *MYB52*, and *MYB93*) that were not identified as direct targets of EIN3 by ChIP-Seq and DAP-Seq (Chang et al., 2013; O’Malley et al., 2016). However, the levels of these transcripts were not significantly altered by ACC in our RT-qPCR data, although there were trends consistent with the patterns observed in the RNA-Seq data (Supplemental Figure 11). This example further validates the ability of this RNA-Seq dataset to identify additional root-specific genes that are dependent upon ETR1 and EIN3 for altered transcript accumulation in response to ACC and/or ethylene treatment.

### The proLRP1::GUS reporter exhibited increased amounts of GUS product in lateral root primordia in the presence of ACC

We were interested in exploring the roles of several of these ETR1-dependent TFs in root development and how they were modulated by ACC. We examined *LRP1* since it had previously been demonstrated to function in lateral root primordium (LRP) development, as *LRP1* overexpression lines had increased numbers of stage I, IV, and V LRP, but reduced numbers of emerged lateral roots (Singh et al., 2020). Since *LRP1* transcripts were induced in whole roots in response to ethylene and ACC treatment (Figure 12A, B), we next asked in which root tissues these transcripts were synthesized. We used the *proLRP1*::GUS transcriptional reporter (Estornell et al., 2018) treated with ACC for five days and visualized GUS product in roots using a stereomicroscope. In 10-day-old seedlings, we found the GUS product accumulated in the central root tissues in both the presence and absence of ACC, beginning at the root elongation zone (Figure 12C). We observed increased GUS activity in the presence of ACC in lateral root primordia at all primordia stages and in emerged lateral roots, with darkest staining evident in the stele layers, consistent with *LRP1* transcripts being synthesized in these tissues and having increased expression in response to ACC treatment.

**Figure 12.**
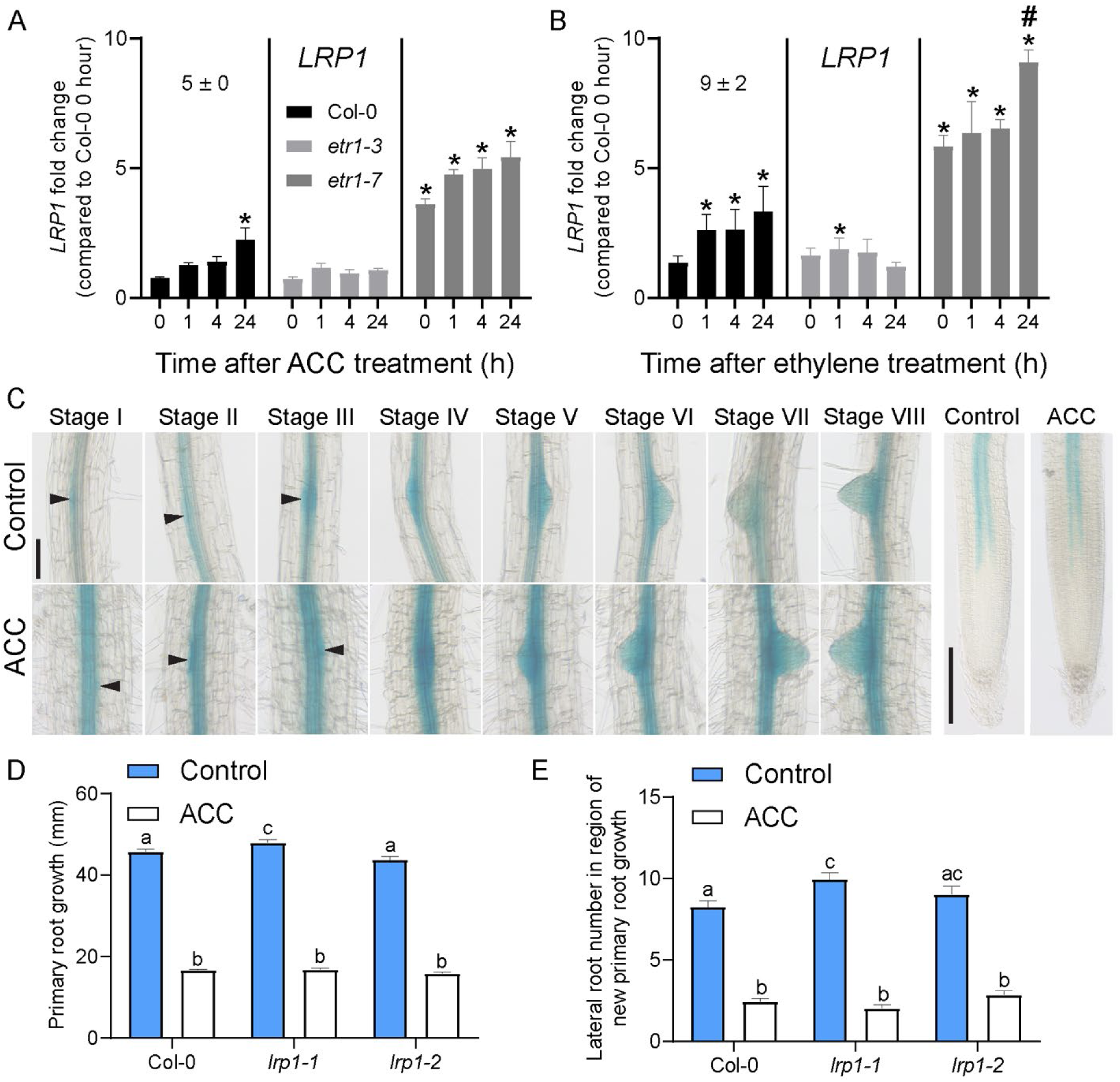
*ProLRP1*::GUS reporter results are consistent with higher LRP1 expression in lateral root primordia in response to ACC treatment. (A, B) *LRP1* normalized transcript abundance was calculated by dividing the TPM of each time point by the average TPM of untreated Col-0. Error bars are SD. (C) Images of lateral root primordia of 10-day-old GUS-stained *proLRP1*::GUS seedlings in the presence and absence of 0.75 µM ACC after five days. Arrows point to stage I-III primordia. Scale bar is 100 µm. Quantification of (D) primary root growth, (E) the number of lateral roots in the region of new root growth in Col-0, *lrp1-1*, and *lrp1-2* in the presence and absence of 0.75 µM ACC after five days. Error bars are SD. Statistical significance is defined as p-value < 0.05 according to Tukey’s multiple comparisons test.

We also quantified the number of lateral roots and the lateral root density in the region of new root growth in the *lrp1-1* and *lrp1-2* mutants in the presence and absence of ACC, with representative images shown in Supplemental Figure 12A. Based on a prior report that examined *LRP1* overexpression lines and identified a 2.5-fold decrease in lateral root density in these lines compared to Col-0 (Singh et al., 2020), we expected an increase in lateral roots in *lrp* mutants. We quantified the number of lateral roots in the region of root formed after transfer to control or ACC media, as we previously demonstrated a negative effect of ACC on lateral root formation in this region (Negi et al., 2008). We found that in time-0 roots, there was a small and significant increase in the number of emerged lateral roots in *lrp1-1* with a non-significant increase in *lrp1-2* (Figure 12E). We found that these small differences were lost in the presence of ACC, which had equivalent inhibition of lateral root formation as Col-0. We also quantified lateral root density in the region of new root growth (5 days of ACC treatment) and found no significant difference in lateral root density in *lrp1-1* and *lrp1-2* in the absence of ACC (Supplemental Figure 12B). The reduction in lateral root density by ACC significantly differed from Col-0 in *lrp1-2* and not *lrp1- 1*, with less inhibition of lateral root formation in *lrp1-2* compared to Col-0. Consistent with the weak accumulation and the absence of changes with ACC of the *proLRP1*::GUS reporter in the primary root tip (Figure 12C), there were subtle changes in primary root elongation in the mutants (Figure 12D). We observed a small but statistically significant increase in primary root growth in *lrp1-1* in the absence of ACC and no significant increase in *lrp1-2* compared to Col-0.

### Several ETR1-dependent TF mutants show subtle differences in root hair initiation, lateral root formation, and root elongation

We screened mutants in a small set of ETR1-dependent TFs for root phenotypes in the presence and absence of ACC. We selected several genes that were also regulated by ACC over a time course of ACC treatment (Harkey et al., 2018). These included *RAP2.6-L*, *ANA058*, *MYB9*, and *MYB52*, some of which were also found to be direct targets of EIN3. To illustrate how these genes responded to ethylene and ACC treatment in Col-0, *etr1-3*, and *etr1-7* in this RNA-Seq study, we reported the fold change in transcript abundance relative to time-0 Col-0 (Supplemental Figure 13). These transcripts were also expressed in various root tissues, including lateral roots, pericycle cells, xylem pole pericycle cells, hair cells, and non-hair cells to varying degrees (Supplemental Figure 14A).

We found that in the region of new root growth, *myb52-1* had significantly fewer lateral roots than Col-0 in the absence of ACC, but it showed wild-type inhibition of lateral root formation in the presence of ACC (Supplemental Figure 14B). The *anac058-1*, *myb52-1*, and *rap2.6l-1* mutants all had lower lateral root densities in this region in the presence of ACC compared to Col-0, suggesting they may function to promote lateral root formation (Supplemental Figure 14C). These results were consistent with *MYB52* and *ANAC058* being down-regulated by ACC, but not with *RAP2.6L* being up-regulated by ACC. The *myb9-1* mutant was also found to have a longer primary root growth phenotype in the absence of ACC compared to Col-0, and no difference was observed in the presence of ACC (Supplemental Figure 14D), suggesting MYB9 may inhibit primary root growth, although this was not consistent with MYB9 being down-regulated by ACC. We also examined the effects of these mutants on root hair number in the absence and presence of four hours of ACC treatment. Only the *myb52-1* mutant had aberrant root hair formation with reduced root hairs compared to Col-0 in both the presence and absence of ACC, suggesting it functions to promote root hair formation (Supplemental Figure 15). These subtle phenotypes were likely due to functional redundancy, as many of the proteins encoded by these transcripts belonged to extensive TF families.

We also compared the growth response of the light-grown seedlings described above to dark-grown seedlings (Supplemental Figure 16). In 10-day-old light-grown seedlings, the *lrp1-1* and *myb9-1* mutants exhibited more primary root growth, yet the ability of ACC to inhibit elongation was like that observed in Col-0, with other mutants having Col-0 levels of root elongation in the absence of treatment (Supplemental Figure 12B; Supplemental Figure 14B). When these same mutants were etiolated on ACC (or untreated) plates for four days, we found that *lrp1-1* had significantly shorter roots than Col-0 at 0.2 µM ACC, while *myb9-1* behaved like Col-0 at all doses of ACC we tested. We also found that *rap2.6l-1*, which did not have a primary root growth phenotype when grown under continuous light, displayed shorter roots than Col-0 at multiple lower doses of ACC (<0.5 µM) after etiolation. However, *lrp1-1* and *rap2.6l-1* did not differ from Col-0 at ACC doses of 0.5 µM and higher. The *myb52-1* mutant displayed longer roots than Col-0 under control conditions at five days of age, although the primary root growth did not differ from Col-0 at 10 days of age when lateral roots were measured. In contrast, etiolated *myb52-1* seedlings displayed longer roots than Col-0 in the absence of ACC treatment but shorter roots than Col-0 in the presence of 0.2 µM ACC. The *anac058-1* mutant was not significantly different from Col-0 at any of the doses of ACC we tested. These differences in light- and dark-grown seedlings suggested that perhaps some of the ETR1-dependent TFs were more important for light-dependent root responses than light-independent ones (Harkey et al., 2019).

## Discussion

The Arabidopsis genome encodes five ethylene receptors that have distinct and overlapping functions in the regulation of plant growth. These receptors negatively regulate the ethylene signaling pathway and are turned off by ethylene binding, inducing downstream changes in transcription. The specific roles of individual receptors in controlling ethylene-regulated development have been examined only in a few tissues (Binder, 2020; Cancel and Larsen, 2002; Harkey et al., 2018; Ma and Dong, 2021). In roots, ETR1 is the ethylene receptor that is most strongly linked to the inhibition of both lateral root formation and primary root elongation and the stimulation of root hair growth (Harkey et al., 2018). To identify the transcripts whose synthesis is regulated by ETR1 to control root development, we identified genome-wide changes in transcript abundance in roots in response to treatment with ethylene or its precursor, ACC, using Col-0 and LOF and GOF *etr1* receptor mutants. We examined the GRNs that are turned on by ethylene and ACC and are dependent on ETR1 signaling, revealing transcripts encoding enzymes of ethylene synthesis and TFs that have the potential to drive transcriptional responses downstream of ethylene.

To identify the transcriptional changes in the presence and absence of ethylene and ACC in Col-0, we treated seedlings with ethylene or ACC and the transcript abundance in roots were quantified at several time points after treatment (0, 1, 4, and 24 hours), and transcript abundance relative to time-0 Col-0 was used to identify DE transcripts and the direction and magnitude change of abundance of these transcripts. Ethylene induced a greater number of DE genes than ACC at all time points after treatment, and the magnitude change of these DE genes was larger (Figure 3). The most likely explanation for this result was that ethylene gas used in our treatments elevated ethylene levels more than this dose of ACC. The majority of changes in transcript abundance in response to either ethylene or ACC treatment were decreases in abundance, especially at the 4- and 24-hour time points. This observation is in alignment with other reports that ACC treatment led to a greater number of transcripts with decreased abundance than increased abundance relative to time-0 controls in Col-0 and in the ethylene-insensitive *ein2-5* mutant (Harkey et al., 2018; Mou et al., 2025).

Several reports have suggested that ACC may act as a signal independently of its conversion to ethylene, especially at high concentrations where ACO-dependent conversion of ACC to ethylene becomes rate limiting (Althiab-Almasaud et al., 2021; Houben and Van de Poel, 2019; Li et al., 2020, 2022; Mou et al., 2020, 2025; Polko and Kieber, 2019; Tsang et al., 2011; Tsuchisaka et al., 2009; Van de Poel, 2020; Van de Poel and Van Der Straeten, 2014; Vanderstraeten et al., 2019; Xu et al., 2008; Yin et al., 2019). At concentrations of 10 µM and above, ACC inhibits rosette size and growth across all tissues in seedlings (Vanderstraeten et al., 2019) and stimulates lateral root initiation and emergence (Mou et al. 2025). However, in roots at lower doses of ACC (less than or equal to 1 µM), both the ethylene-insensitive *ein2-1* and *ein2-5* mutants do not respond to treatment with 1 µM ACC (Harkey et al., 2018; Ivanchenko et al., 2008; Muday et al., 2012; Negi et al., 2008; Růžička et al., 2007), suggesting that ACC is either largely converted to ethylene at this dose or that in these tissues under these growth conditions ACC predominantly acts via its conversion to ethylene. Our finding that most of the transcripts that were DE in response to this low dose of ACC were also DE in response to ethylene is consistent with these possibilities. Yet the presence of transcripts in this dataset whose abundance changed after regulated by ACC-treatment, but not ethylene-treatment, even though ethylene generally resulted in more robust transcript abundance changes, is consistent with ACC having a direct effect on transcription.

A recent article described a mechanism by which ACC-regulated transcripts may result from direct ACC signaling to TFs rather than signaling after its conversion to ethylene. ACC has been shown to regulate WOX5 and LBD18 to control meristem growth and lateral root development, respectively, independent of ethylene signaling (Mou et al., 2025). The majority of the positively ACC-regulated DE transcripts identified in their udataset when ethylene- insensitive *ein2-5* seedlings were treated with ACC were also DE in response to ACC in our dataset, although not the downregulated genes, consistent with conserved transcriptional responses between our datasets.

We found that several transcripts regulated by only ACC included those encoding MAP kinases, enzymes that remodel the cell wall, and proteins involved in root hair development, as well as ACO enzymes whose regulation we characterized. We found that ACC down-regulated *ACS7*, which might suggest a negative feedback of ACC synthesis in response to elevated levels of ACC, while *ACO* transcripts increase to facilitate conversion to ethylene.

An important goal of these experiments was to take the large number (8,323) of DE genes in response to ethylene and/or ACC (Figure 4) and refine this dataset by identifying the genes whose regulation of depended on ETR1. We identified ETR1-regulated genes using two *etr1* mutant alleles: *etr1-3*, which is a GOF receptor mutant with a constitutively active receptor that turns off ethylene signaling, resulting in an ethylene-insensitive phenotype, and *etr1-7*, which is a LOF receptor mutant that has an inactive receptor, resulting in constitutive ethylene signaling. The combination of GOF and LOF receptor mutants served as a powerful approach to identify ETR1-dependent transcriptional responses. This is clear from the earliest steps of the analysis; comparison of all samples by a PCA plot, Pearson’s correlation, and Euclidean distance plots led to the separation of *etr1-7* samples from Col-0 and *etr1-3* on the PC1 axis (Figure 2A).

We used a strict criterion to define the ETR1-dependent genes, requiring genes were DE with either ethylene or ACC, significantly altered in abundance in the *etr1-7* allele, with constitutive ethylene response, and transcripts were not significantly changed by ethylene or ACC in the ethylene insensitive *etr1-3* mutant. The pattern of response evident in Figure 6 shows enhanced transcript abundance in *etr1-7* that is not changed by either ethylene or ACC treatment, with these transcripts having muted response to both hormones. The strict ETR1-dependent group was enriched in transcripts annotated to function in “ethylene mediated signaling pathway.” We also compared our ETR1 dependent genes to a conserved group of ethylene- responsive genes from a meta-analysis of three root ACC and ethylene response datasets (Harkey et al. 2019) finding 31 transcripts that were ETR dependent in the overlap of these groups.

An unexpected finding in this study was that there were nearly as many transcripts that responded in an ETR1-independent fashion as those that were ETR1-dependent (Figure 7). Transcripts were defined as ETR1-independent if they responded to ACC or ethylene treatment in all three genotypes. If the abundance of these transcripts increased and decreased in *etr1-7* and *etr1-3*, respectively, and changed with hormone treatment, this might suggest two receptors control their activity, with hormone-dependent changes being mediated by a second receptor. In a prior study we found ETR1 to be essential in some root developmental responses and identified roles for other ethylene receptors in other responses (Harkey et al., 2018). ACC effects on lateral root development were completely lost in the ethylene-insensitive mutant *etr1-3* and not induced by addition of ACC in the constitutively signaling mutant *etr1-7*, suggesting inhibition of lateral root formation by ethylene signaling had strong dependence on ETR1. In contrast, inhibition of primary root elongation and stimulation of root hair growth by ACC were less dependent on ETR1 as they showed attenuated responses to ACC in *etr1-7* (Harkey et al., 2018), suggesting a role of other ethylene receptors in these root processes. Perhaps some of the transcripts which were ETR1 independent were involved in root elongation or root hair initiation. Indeed, when we examined genes that were ETR1-independent by the strictest criteria, they were enriched in the annotation “root hair cell differentiation” and several other related annotations.

We had identified a small subset of transcripts that were regulated only by ACC in Col-0, suggesting responses that may be driven by ACC signaling independently of its conversion to ethylene. Therefore, we were interested in further analyzing the transcripts that encode ACC oxidases, which convert ACC to ethylene, as this might provide insight into understanding how ACC may regulate its conversion to ethylene.

In our dataset, we found that *ACO* transcripts were themselves regulated by ACC and ethylene in a positive feed forward loop, where application of ethylene and/or ACC induced transcript accumulation of all *ACO*s in at least one time point after treatment (Figure 8). We found that *ACO2* and *ACO3* met our strict criteria for ETR1 dependence in response to ethylene but not ACC. This suggests that ACC may be able to regulate transcripts in ways above and beyond the function of ETR1, consistent with the possibility of an ACC-independent signaling pathway. However, for *ACO2*, perhaps the most puzzling of the *ACO*s, ACC-mediated changes in transcript abundance were completely lost in *ein3eil1*, suggesting that ACC activates the canonical ethylene signaling pathway to control *ACO2* levels. It is not clear how ACC both activates the canonical ethylene signaling pathway and induces changes in the *etr1* mutants that were not induced by ethylene. Perhaps ACC can work through a non-canonical ethylene signaling pathway or directly bind to a different ethylene receptor, as this might be required when ACC levels become too high, and the plant needs a mechanism to rapidly convert ACC to ethylene. There is evidence that glucose can signal to EIN2 and subsequently EIN3/EIL1 through the target of rapamycin pathway (Fu et al., 2021), so it is possible that ACC could activate this or a similar pathway resulting in transcriptional changes that are not dependent on ETR1 but require the EIN3/EIL1 machinery.

To provide more information on the localization of changes in *ACO* transcript abundance, we examined the root cell-type-specific localization of *ACO* transcripts in untreated roots using publicly available transcriptome data and *in planta* using ACO promoter-driven fluorescent protein reporters to inform our understanding of the tissues that express the machinery to convert ACC to ethylene under our treatment conditions (Figures 9 and 10). We found that each *ACO* had a distinct tissue expression pattern, and many of these were enhanced in the presence of ACC treatment in 10-day-old seedlings. These findings were consistent with *ACO*s being expressed in root hairs, lateral roots, pericycle cells, and the root elongation zone where their protein products convert ACC to ethylene to signal root responses in these tissues. The most striking observation was that *ACO5* transcripts were the only *ACO* accumulating to high levels in hair cells prior to and after root hairs initiated, and this root hair expression was evident in ACC- treated seedlings that formed more and longer root hairs, consistent with a role for ACO5 in driving root hair elongation. The other *pACO*::GFP reporters were not detected or had low fluorescence in root hairs.

We were interested in mapping the GRN downstream of ETR1 and EIN3/EIL1 by examining transcripts predicted to encode TFs using the Agris database (Davuluri et al., 2003). We identified 60 TF-encoding transcripts in the ETR1-dependent list of genes and found some that were rapidly up-regulated and a greater number that were down-regulated by ACC and ethylene treatment with a range of different kinetic responses to these treatments (Figure 11). We also generated a network using PaLD, which groups transcripts together with the most similar temporal responses. This network allows us to visualize connections among transcripts with the most similar responses to ACC and ethylene across time and genotypes and to partition transcripts into groups of like responding genes. We found that the up-regulated genes clustered into two groups, one consisting of transcripts with rapid and strong induction in response to treatment, and another consisting of transcripts with more muted up-regulatory patterns, and both groups were enriched in direct targets of EIN3. In contrast, the down-regulated genes clustered into four groups containing two or more transcripts with distinct expression patterns, and few of the transcripts in these groups were direct targets of EIN3. These results raise the interesting question of what is the regulatory mechanism by which so many of these transcripts have decreased abundance in response to ACC and ethylene treatment.

We asked whether three of the TF-encoding genes that were found to be EIN3 targets in DAP-Seq or ChIP-Seq datasets (Chang et al. 2013; O’Malley et al. 2016) lost their ACC-regulated expression in an *ein3eil1* mutant, focusing on *ROOT HAIR DEFECTIVE 6-LIKE 5* (*RSL5*), *RELATED TO AP2 6L (RAP2.L)*, and *MYB9* (Figure 11). We found that the ACC-mediated increase in *RSL5* and *RAP2.6L* transcripts and the decrease in *MYB9* transcripts were both lost in *ein3eil1*. It was previously reported that an *rsl5* mutant exhibited normal root hair development in the absence of treatment (Pires et al. 2013), which is not surprising due to the low levels of transcript abundance (2 TPM) in time-0 roots, suggesting that RSL5 might function in ACC- and ethylene-stimulated root hair formation. An interesting question that arises from this work is how ETR1 regulates genes that are not regulated by EIN3/EIL1. It is possible that ETR1-dependent genes could be regulated by the non-canonical AHP and ARR families (Binder et al. 2020), as previous work demonstrated direct interaction and phosphorylation between ETR1 and AHP1 in vitro (Scharein and Groth, 2011).

ETR1 is important in ethylene’s ability to inhibit lateral root formation in the region of new root growth formed after exposure to ACC. *LRP1* was identified as ETR-dependent in our dataset; this gene encodes a TF that has previously been shown to function in inhibiting lateral root emergence downstream of ARF7/ARF19, LATERAL ROOT PRIMORDIA 1 (LRP1) (Singh et al., 2020). LRP1 overexpression lines had fewer emerged lateral roots, suggesting that LRP1 functions to inhibit lateral root emergence. By examining a *proLRP1*::GUS reporter, we found that *LRP1* expression was increased by ACC in lateral root primordia at all stages of primordia development, and in emerged lateral root tips and in the innermost root cells of primary and lateral roots (Figure 12). However, we examined two *lrp1* mutant alleles in the presence and absence of ACC we observed that one mutant had reduced inhibition of lateral root formation in response to ACC, while the other allele did not, which is inconsistent with the wild-type gene functioning to control lateral root development under our growth conditions.

We examined mutants in several other TFs that were selected based on their transcriptional responses to ACC in both this dataset and our prior ACC time course dataset (Harkey et al., 2018), including ANAC058, MYB9, MYB52, and RAP2.6L (Supplemental Figures 15, 16, and 17). We found subtle but statistically significant root phenotypes in these mutants, with *anac058-1*, *myb52-1*, and *rap2.6l-1* having reduced lateral root densities in the presence of ACC compared to Col-0, and *myb9-1* having increased primary root growth compared to Col-0 in the absence of ACC. We also found that *myb52-1* had fewer root hairs than Col-0 in both the presence and absence of ACC. We also examined these mutants for triple response phenotypes in the presence and absence of several concentrations of ACC, finding differences in phenotypes between seedlings grown under continuous light conditions and those grown in the dark for four days prior to exposure to light. These findings suggested that some of these TFs could be involved in the light-dependent pathway downstream of ethylene (Harkey et al., 2019).

This study provides insight into the ETR1-mediated GRN controlling root growth and development, narrowing the previously identified genes that are regulated by ethylene and ACC into a group of transcripts that are regulated by ETR1 and/or EIN3. We examined the transcriptional regulation of genes encoding ACC oxidases to understand this final step in ethylene synthesis, which controls the balance of accumulation of the ACC precursor and the ethylene product. We find that the *ACO* transcript abundance is regulated more strongly by ACC than ethylene consistent with feed forward synthesis of ethylene when ACC accumulates, which hints at an ethylene-independent role in the regulation of this gene family. This study also provides a valuable dataset to allow researchers to further dissect the role of ETR1 in controlling root responses to changes in both ethylene and the ethylene precursor, ACC.

## Methods

### Genotypes and plant growth conditions

All mutants used in this study were in the Col-0 background. Both the *etr1-3* and *etr1-7* alleles have been described previously (Guzmán and Ecker, 1990; Harkey et al., 2018; Hua and Meyerowitz, 1998). The *etr1-7* (AT1G66340) mutant was obtained from Elliot Meyerowitz (Hua and Meyerowitz, 1998). T-DNA insertion mutants were obtained from the Arabidopsis Biological Resource Center for AT3G18400 (SALK_049205C), AT5G12330 (SAIL_402_G06/CS873836 and SALK_201247C), AT5G16770 (SALK_149765C), AT1G17950 (SALK_138624C), and AT5G13330 (SALK_051006C); see Supplemental Table 5 for more information on these lines. The *pACO*::GFP-GUS reporters were prepared by Dr. John Vaughan-Hirsh and shared by Dr. Bram Van de Poel (Houben et el. 2024). The *proLRP1*::GUS reporter was obtained from ABRC (Estornell et al., 2018).

Plants were grown on 1× Murashige and Skoog medium (Caisson Laboratories), pH 5.6, Murashige and Skoog vitamins, and 0.8% agar, buffered with 0.05% MES (Sigma), and supplemented with 1% sucrose. After stratification for 72 hours at 4°C, plants were grown under 100-130 µmol m^−2^ s^−1^ continuous cool-white light.

For selecting a dose of ethylene for the RNA-Seq, Col-0 seedlings were treated on day five after germination. For ACC treatment, the seedlings were transferred to a growth medium containing 0 or 0.75 µM ACC for 24 hours before imaging. For ethylene treatment, plates were placed in clear treatment tanks with constant flow-through of the indicated concentration of ethylene for 24 hours before imaging.

### Method for selecting time points for RNA-Seq analysis using a previous microarray study

We developed an analysis approach to identify the subset of time points that contained the most unique (i.e., not found in other time points) DE genes in response to ACC treatment using our previously published time-course microarray study with ACC treated Col-0 roots (Harkey et al. 2018). In that prior study, we examined transcriptional responses to 1 µM ACC after 0, 0.5, 1, 2, 4, 8, 12, and 24 hours of treatment, which overlaid the timeline for ACC-induced root hair initiation. To inform the selection of time points for the present study, we first analyzed the prior microarray data to find DE genes for each individual time point compared to time 0. We then performed a pairwise comparison of time points to determine the number of overlapping DE genes between each time point (Supplemental Figure 2B) and the percent of DE genes in a time point that overlapped with another time point (Supplemental Figure 2A). These results revealed that treatment times of 1, 4, and 24 hours contained the most unique sets of transcripts while also overlapping with the timing of root hair formation to capture early, middle, and late developmental responses. Therefore, we chose to perform RNA-Seq using samples treated with ACC or ethylene at three time points: 1 hour, to monitor rapid transcriptional responses that precede developmental changes; 4 hours, where the most unique transcripts are DE and root hair initiation and primary root elongation are increasing and decreasing, respectively, in response to treatment; and 24 hours to identify transcripts linked to ethylene- modulated lateral root regulation.

Microarray data (Harkey et al., 2018) was analyzed using limma (Ritchie et al., 2015). Probes were first filtered for interquartile range (IQR), with anything less than 0.1 excluded from further analysis. Time points were then compared to their time-matched controls; p-values were adjusted using a Benjamini-Hochberg correction. Genes with an absolute log_2_FC greater than 0.5 and an adjusted p-value less than 0.05 were considered to be DE at that time point. The overlap matrix was created by comparing the DE genes across the time points in a pair-wise manner.

Supplemental Figure 1A graphically illustrates the results of this analysis. For each column, the percentage of DE genes at that time point (column) that are also DE in the time point to which it was compared (row) is shown. The number of genes in each comparison that was used to calculate the percentages can be seen in Supplemental Figure 1B. For example, the 2- hour time point showed considerable overlap with the four-hour time point; 87% of all 2-hour DE genes were also DE at 4 hours. The reverse comparison of the four-hour genes that were also DE at 2 hours yielded only 32% of the four-hour genes that were DE at two hours because there were more DE genes in this 4-hour sample. Therefore, the four-hour time point identified nearly all the same DE genes as the 2-hour time point, as well as many more unique genes, so we selected the 4-hour time point over the 2-hour for this RNA-Seq analysis. We also examined the overlaps of the 4- and 8-hour time points and the 12- and 24-hour time points, finding more unique DE transcripts in the 8- and 24-hour time points for each of these overlaps. However, as root hairs had already begun to initiate by 4 hours (Harkey et al., 2018), and the ethylene-dependent inhibition of lateral root initiation had begun by 24 hours, these time points also represented important developmental milestones. Ultimately, this analysis informed our choice to do the present experiment with time points 0, 1, 4, and 24 hours.

### ACC and ethylene treatment of seedlings for RNA isolation for sequencing

Plants were grown on media as described above on top of a nylon filter (03-100/32; Sefar Filtration) pressed against the plate, as described previously (Harkey et al., 2018; Levesque et al., 2006). Approximately 100 sterilized seeds were placed on each filter; 2 plates were combined for each biological replicate. ACC-treated samples were grown and harvested at Wake Forest University, and ethylene-treated samples were grown and harvested at the University of Tennessee Knoxville utilizing a gas flow-through system to treat with constant levels of ethylene gas. The same researchers prepared both sets of samples, and identical reagents and light treatments were used in both laboratories. Time-0 controls were prepared in both locations to account for any location-specific effects on baseline expression.

Plants were treated on day five after germination. For time-0 samples, the nylon filter containing five-day-old plants was transferred to a new control medium and root tissue was immediately harvested. For ACC treatment, the plants on nylon were transferred to growth medium with 1 µM ACC for the given treatment time (1, 4, and 24 h) and then root tissue was harvested. For ethylene treatment, the nylon was transferred to new control medium, and plants were placed in clear treatment tanks with constant flow-through of 0.3 ppm ethylene gas for the given treatment time (1, 4, and 24 h) and then root tissue was harvested. At time of harvesting, roots were cut from seedlings and flash frozen in liquid nitrogen. Frozen samples were ground in liquid nitrogen, and RNA isolation was performed according to the Qiagen plant RNeasy kit protocol, with the addition of the Qiagen RNase-free DNase treatment (Qiagen). After RNA isolation, samples were quantified by A260 using a Nanodrop spectrophotometer (Nanodrop Technologies). Each sample yielded at least 3 µg, and on average 15 µg, of RNA. One Col-0 sample which had a 4-hour ethylene treatment had an RNA integrity number (RIN) less than 6 and was not sequenced. Sequencing was performed by GENEWIZ, LLC using the Illumina HiSeq platform.

### RNA-Seq preprocessing and quality control

FastQC v0.11.8 (Wingett and Andrews, 2018) and MultiQC v1.7 (Ewels et al., 2016) were used for assessing read quality, which identified adaptor contamination and a bimodal GC Content distribution. Adaptors were trimmed from reads using CutAdapt v1.18 (Martin, 2011), and reads trimmed to less than 25 bp were removed. Trimmed reads were then processed with the BBMap tool BBDuk (Bushnell 2014) to identify and remove rRNA contamination. Running FastQC/MultiQC over the trimmed and filtered reads revealed the identified quality issues had been mitigated. Read quantification was performed directly on the processed fastq files with the Salmon v0.12.0 (Patro et al., 2017) “quant” algorithm in mapping mode using the “-- validateMapping” flag, setting the library type to “IU”, and using the TAIR10 reference transcriptome (NCBI ID: GCF_000001735.3). Counts were imported into R v3.6.0 using the tximport R package v1.12.3 (Soneson et al., 2016) and summarized to the gene level.

### Differential expression and visualization

For visualization and sample comparisons, raw read counts were normalized using the Variant Stabilizing Transformation method from the R package DESeq2 v1.24.0 (Love et al., 2014). Sample-wise comparison heatmaps were generated using Pearson’s correlation and Euclidean distance as the distance metrics, respectively. PCA analyses were performed by DESeq2’s plotPCA method using the top 1000 most variable genes for all genotypes together, and on a per-genotype basis.

All sample-wise comparisons were performed using the DESeq2 package and ashr v2.2- 47 (Stephens et al., 2020). Density plots were generated using ggplot2 v3.3.0 (Wickham, 2016), and heatmaps of sample-wise comparisons were generated using ComplexHeatmap v.2.3.4 (Gu et al., 2016). For DE transcripts used in heatmaps and GO analyses, we considered a transcript to be DE if it had a p-value < 0.01, |logFC| > 0.5, and baseMean > 10.

### Identification of outlier and pooling of control samples

We performed ACC treatments at Wake Forest University and ethylene treatments at the University of Tennessee. Therefore, to determine if these samples were similar, we first compared the controls from each location for each genotype (Supplemental Table 1). Since the controls for Col-0 and *etr1-3* were well matched between the two locations, and the PCA plots suggested differences in *etr1-7* were not location dependent (Figure 2D), we combined the six control (0-hour) samples for each genotype, or the five control samples for Col-0, excluding the outlier sample described above, to use as the baseline for comparisons for the remaining analyses.

To determine if all the Col-0 control (time-0) samples could be pooled for downstream analyses, we generated a PCA plot using only the Col-0 samples (Supplemental Figure 3). This plot revealed that one sample (in the ethylene time-0 treatment group) did not cluster with the other control samples. This sample showed substantial separation from the rest along the PC1 axis, which represents over 50% of the variance across all the Col-0 samples. This outlier sample was found to have far lower RNA yield than the other 71 samples, so we removed it from downstream analyses. We generated a new PCA plot without this sample, revealing much greater consistency.

To further validate this method, we performed two DE analyses to examine batch effects on samples collected in the two locations, one with all six Col-0 control samples included (three from each location) and one with the outlier sample removed. When all six samples were included, 693 genes were found to be DE (adjusted p-value < 0.01, |log_2_FC| > 0.5) between the ACC and ethylene controls performed at these two locations (Supplemental Table 1). When the outlier sample identified by the PCA plot was removed from the analysis, only 34 genes were found to be DE between the Col-0 samples (Supplemental Table 1). This removal also affected the model generated by DESeq2, resulting in subtle effects on the *etr1-3* and *etr1-7* comparisons (Supplemental Table 1). In the model with the outlier Col-0 removed, 121 and 763 genes were DE between the controls compared at each location in *etr1-3* and *etr1-7*, respectively (Supplemental Table 1). The time-0 *etr1-7* samples had more DE genes between locations than had been expected, but there were two controls that were quite different than other samples (as evident in the PCA analysis in Figure 2D), which suggested that this is a character of two samples rather than a difference in samples because of location. We ultimately decided to combine the control samples for each genotype from both locations.

### PaLD analysis

A newly developed tool called Partitioned Local Depths (PaLD) was used to generate networks of the 553 ETR1-dependent transcripts and the 60 ETR1-dependent transcription factors, with application of this algorithm to gene expression and time course datasets described previously (Berenhaut et al., 2022; Khoury et al., 2024a). We sorted genes based on their transcript abundance patterns in response to ACC and ethylene over time in Col-0, *etr1-3*, and *etr1-7*. We used the log_2_FC values relative to time-0 Col-0 as input for the PaLD analysis. We further divided the networks using the Louvain method for community detection (Blondel et al., 2008). Relevant R code can be found at https://github.com/moorekatherine/pald.

### RT-qPCR

RNA was extracted from roots using the RNeasy Plant Mini Kit (Qiagen) according to the manufacturer’s protocol, with buffer RLT selected as the extraction buffer for roots. RNA with A260/230 values less than 1.5 were re-isolated using isopropanol. DNA was removed using the RapidOut DNA Removal Kit (ThermoScientific) according to the manufacturer’s instructions. Complementary DNA was synthesized using the Superscript III First-Strand Synthesis System for RT-PCR (Invitrogen) with RNA template primed with random hexamers according to the manufacturer’s instructions. PowerUp SYBR Green Master Mix (Applied Biosystems) was used for quantitative PCR on a QuantStudio 3 Real-Time PCR System (Applied Biosystems). Primer efficiency and transcript abundance values were calculated using the standard curve method and primer sequence is reported in Supplemental Table 1.

### Fluorescence and brightfield imaging

For high resolution imaging of all *pACO*::GFP-GUS reporters, a Zeiss LSM 880 microscope was used. Roots were first stained with propidium iodide (PI) diluted in water to a final concentration of 0.01 mg/mL for 2 min before being washed briefly in DI water and mounted on slides in DI water with #1.5 coverslips. Confocal images were acquired with a 488 nm excitation laser for imaging GFP and the 561 nm laser for imaging PI with emission ranges of 490-552 nm and 56-696 nm, respectively. To minimize light scatter and improve image resolution, the pinholes for imaging GFP and PI were adjusted to slightly less than 1 Airy Unit. Images were taken with a Plan-Apo 20x/0.8NA objective and field size of 2048 x 2048 pixels. For Z-stacks, pixel dwell time was 0.26 µs and line averaging was set to 4. Z-stacks were obtained with 2.9 µm intervals between optical slices. For single-slice, high-resolution images, pixel dwell time was 1.02 µs and line averaging was set to 8.

For epifluorescence imaging of the *pACO5*::GFP-GUS reporter, a Zeiss Axio Zoom.V16 equipped with a PlanNeoFluar Z 1.0x (0.25NA) objective was used. An excitation range of 430-495 nm and emission range of 495-575 nm was employed with a Zeiss 38HE fluorescence filter cube. Images were captured with an Axiocam 506 monochrome camera with an exposure time of around 500 ms.

For imaging *proLRP1*::GUS, the Zeiss Axio Zoom.V16 equipped with a PlanNeoFluar Z 2.3x (0.75NA) objective was used. Images were captured with an AxioCam HR R3 color camera with an exposure time of around 10 ms.

### GUS staining procedure

Fresh X-Gluc (GoldBio) was dissolved on ice in N,N-Dimethylformamide immediately before incubating samples. For GUS staining, ten-day-old seedlings were vacuum infiltrated with GUS buffer containing X-Gluc (100 mM phosphate buffer pH 7, 0.5 mM K_4_Fe(CN)_6_, 0.5 mM, K_3_Fe(CN)_6_, 0.5 mM, 10 mM EDTA, 0.1% Triton X-100, and 2 mM X-Gluc) for five minutes, then incubated in the buffer and X-Gluc mixture for 1.5 hours at 37°C. Following product formation, seedlings were washed in 70% EtOH for 10 min. then fixed overnight in 3:1 EtOH:acetic acid at 4°C in the dark prior to imaging.

### PCR genotyping

The primers used for genotyping are reported in Supplemental Table 5. Young leaf tissue was homogenized in a microcentrifuge tube using a micropestle that was dipped in liquid nitrogen. Edwards Solution (Edwards et al., 1991) was added to the homogenized leaf tissue and samples were incubated at room temperature for at least one hour before being spun down. Supernatant was carefully transferred to a new tube containing an equal volume of isopropanol.

Samples were washed with EtOH and air dried in a sterile hood for approximately 30 minutes before being resuspended in molecular biology grade H_2_O. Genotype was determined using wild-type primers that flanked the T-DNA insert (the wild-type reaction) to amplify the gene, and right border primers with T-DNA primers (the T-DNA reaction) to amplify the T-DNA insert (O’Malley et al., 2015), with primer sequences listed in Supplemental Table 5. DreamTaq DNA Polymerase (ThermoFisher) was used according to the manufacturer’s instructions.

### Root development analysis

For analysis of root hair and lateral root phenotypes, five-day-old seedlings were transferred to plates containing 0 or 0.75 µM ACC for 4 hours or 5 days before imaging root hairs and lateral roots, respectively. For the root length analysis of dark grown seedlings, seeds were germinated in the dark for 4 days on untreated plates or plates containing 0.1, 0.2, 0.5, 1 or 2 µM ACC, then root lengths were measured. Statistical tests (two-way ANOVA and multiple comparisons tests) were performed in GraphPad Prism 8.0.2. Outliers were identified using ROUT (Q = 1%).

## Acknowledgements

This work was supported by the National Science Foundation Systems and Synthetic Biology (MCB-1716279 to GKM) and the Center for Molecular Signaling Graduate Research Fellowship to MGW and a WFU Provost’s Pilot Research grant (to GKM and MGW). We appreciate the help and mentorship of Megan Gerber, particularly the comparison of transcriptomic datasets.

## Author Contributions

Maleana White, Alexandria Harkey, and Amy Olex helped design the research, performed and analyzed data and wrote and edited the manuscript. Joelle Muhlemann, Nathan Pfeffer, and Maarten Houben performed experiments, analyzed data, and edited the manuscript. Brad Binder helped design and performed experiments and edited the manuscript. Gloria Muday designed experiments and wrote and edited the manuscript.

