## Supplemental Tables and Figures for "Ethylene Receptor Gain- and Loss-of-function Mutants Reveal an ETR1-dependent Transcriptional Network in Roots"

**Supplemental Materials**

**Supplemental File 1.** This file includes a list of genes that were DE in response to ethylene and/or ACC in Col-0 at each time point, a summary of ETR1 dependence, a list of the ETR1-dependent, -independent, and complex DE genes, and a list of the DE genes in the genotype comparisons at baseline. All DE genes are listed with their $log_2FC$ values (relative to Col-0 control samples or other genotype in the genotype comparisons) and p-values.

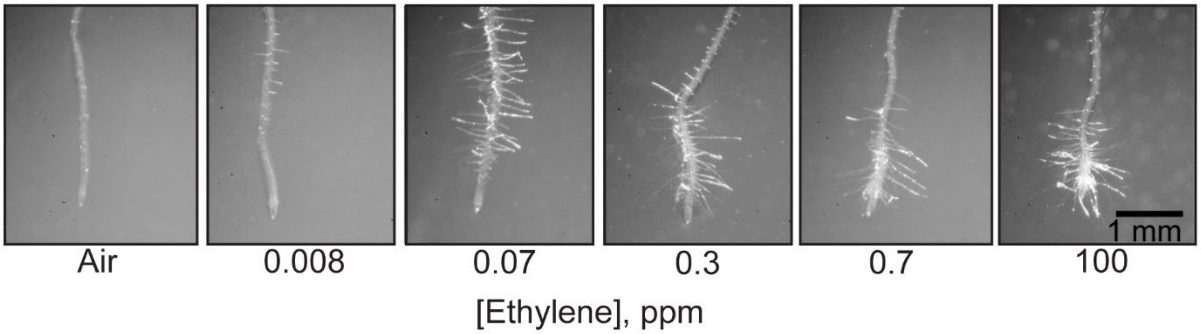

**Supplemental Figure 1. Root hair formation in response to increasing concentrations of ethylene gas.** Images of the root tips of six-day-old seedlings were captured 24 hours after treatment with ethylene gas at the indicated concentrations.

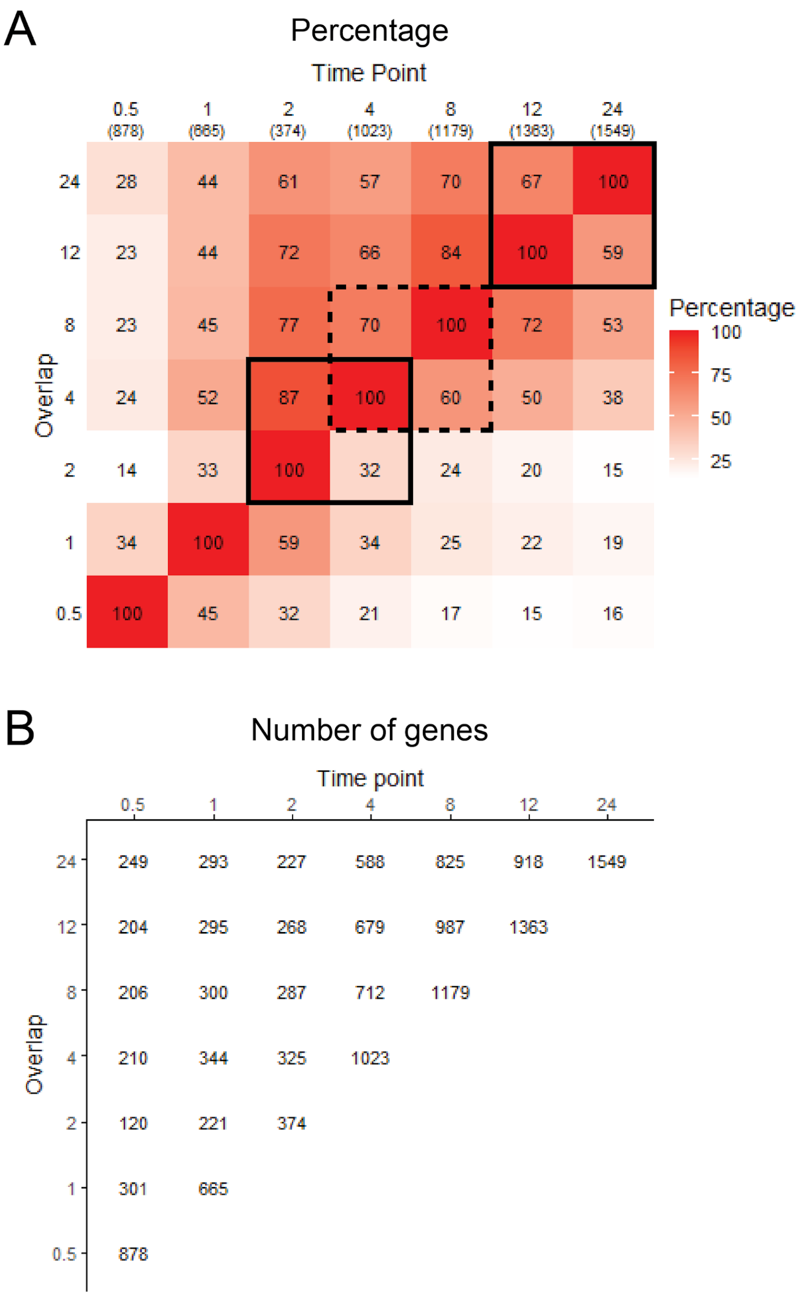

**Supplemental Figure 2. Pairwise comparison of the DE genes at each time point in the ACC microarray dataset** (Harkey et al., 2018)**.** (A) The percentage of DE genes in one time point (column) which were also DE in the overlap time point (row). The total number of DE genes in each time point is shown in parentheses under the column time point labels. Comparisons highlighted in the text are boxed in black solid or dashed lines. (B) The number of DE genes in one time point (column) which were also DE in the overlap time point (row).

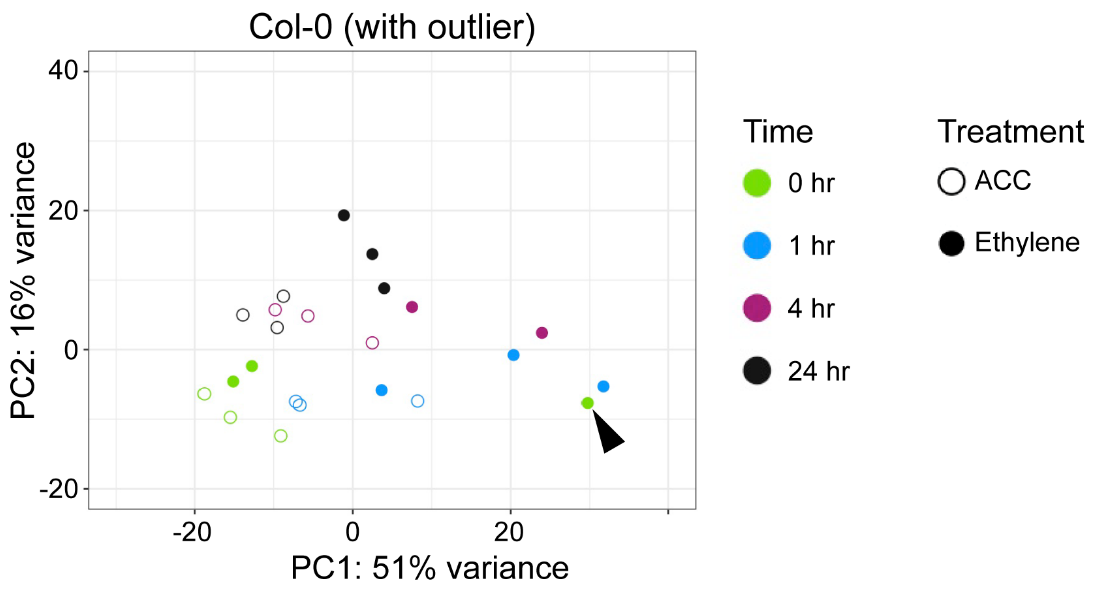

**Supplemental Figure 3. Principal component analysis was used in the decision to remove Col-0 outlier.** PCA plot of Col-0 samples, with the outlier sample included.

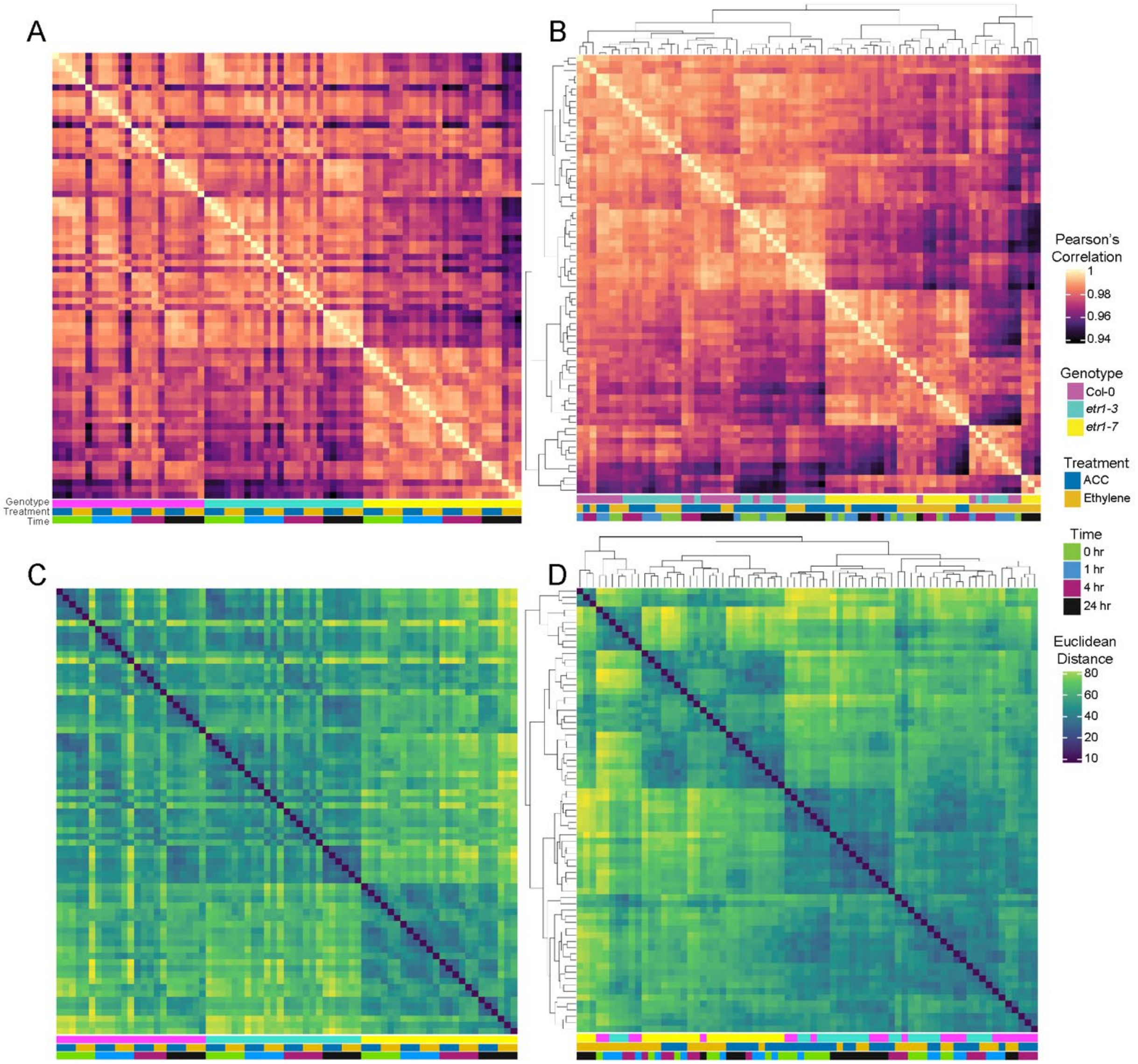

**Supplemental Figure 4. Large-scale patterns of data from all samples.** Samples were compared in a pair-wise manner using Pearson's correlation (A & B) or Euclidean distance (C & D), then sorted by genotype and treatment (A and C), which highlights samples that differed from other replicates from the same genotypes and treatment times, or clustered based on greatest similarity between samples (B & D).

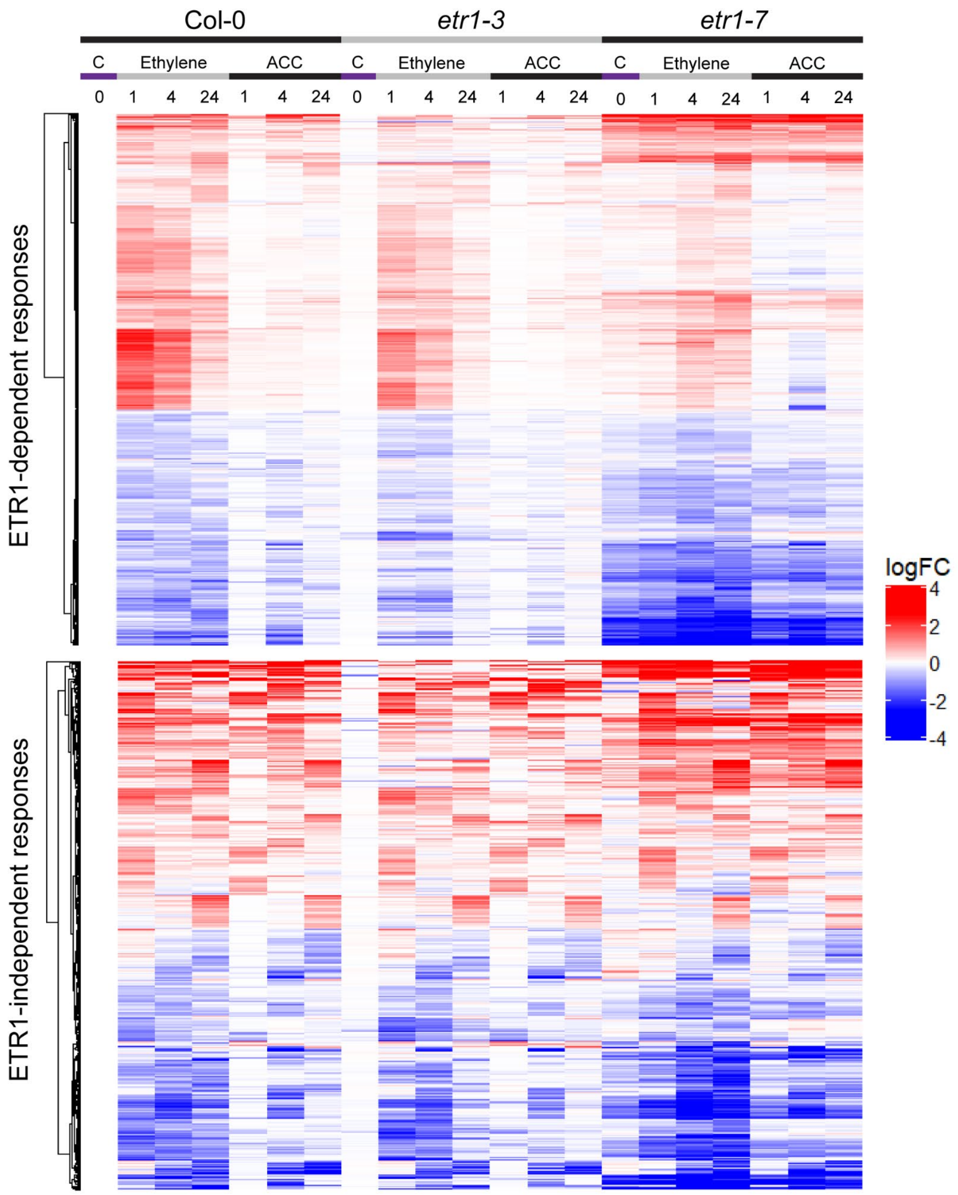

**Supplemental Figure 5. Transcripts identified by less stringent criteria for ETR1 dependence or independence.** For this analysis, ETR1-dependent transcripts were required to respond to treatment in Col-0, and not respond in the mutants; ETR1-independent transcripts were required to respond to treatment in all three genotypes. Both sets of criteria were evaluated based on comparison of treatment samples to the untreated control samples for the same genotype, regardless of the transcript abundance in that control sample.

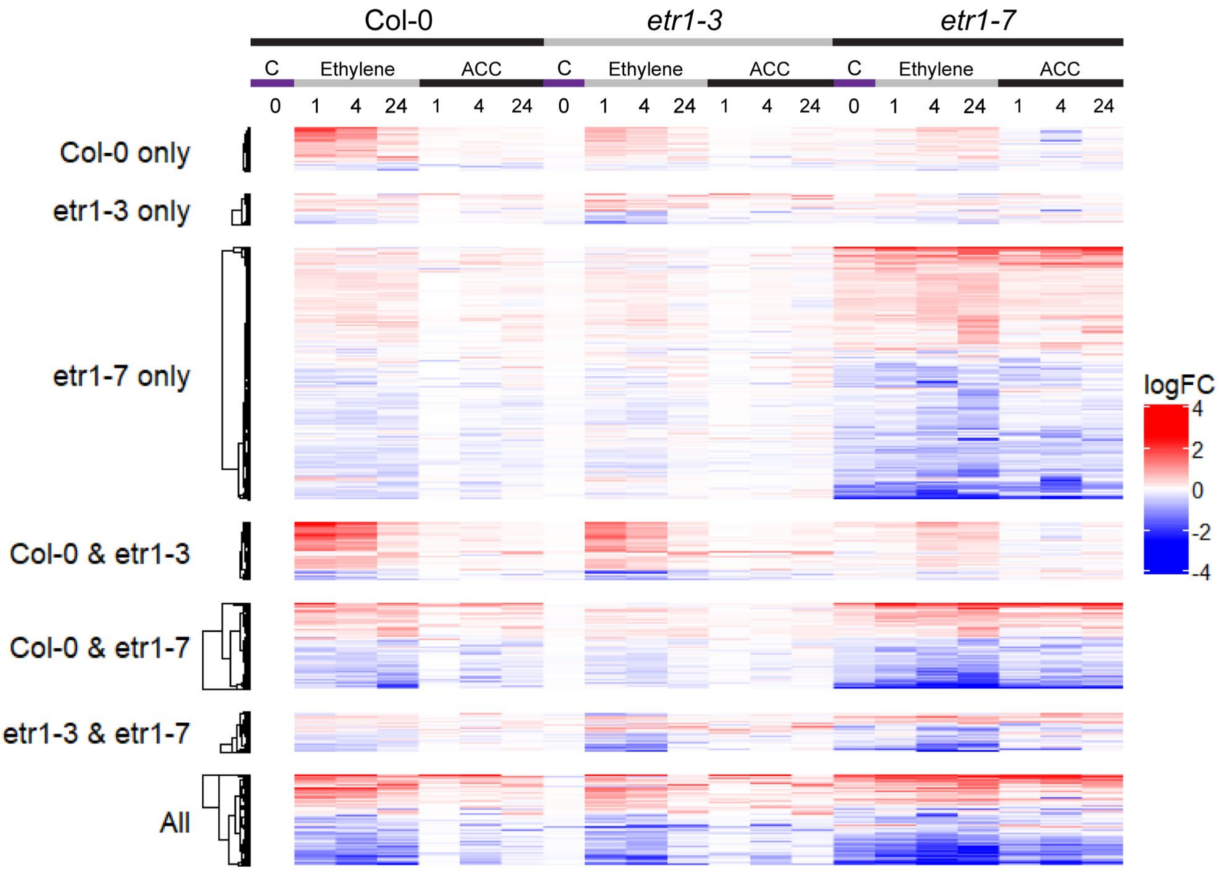

**Supplemental Figure 6. Transcripts with complex responses, which responded to ACC or ethylene, but were not consistently regulated by ETR1.** Genes with complex responses to ethylene and/or ACC which did not qualify as ETR1-dependent or -independent. Genes were split into groups based on which genotypes had at least one DE sample, based on $|\log_2 FC| > 0.5$ and adjusted p-value < 0.01. $\text{Log}_2$FC was reported relative to untreated Col-0.

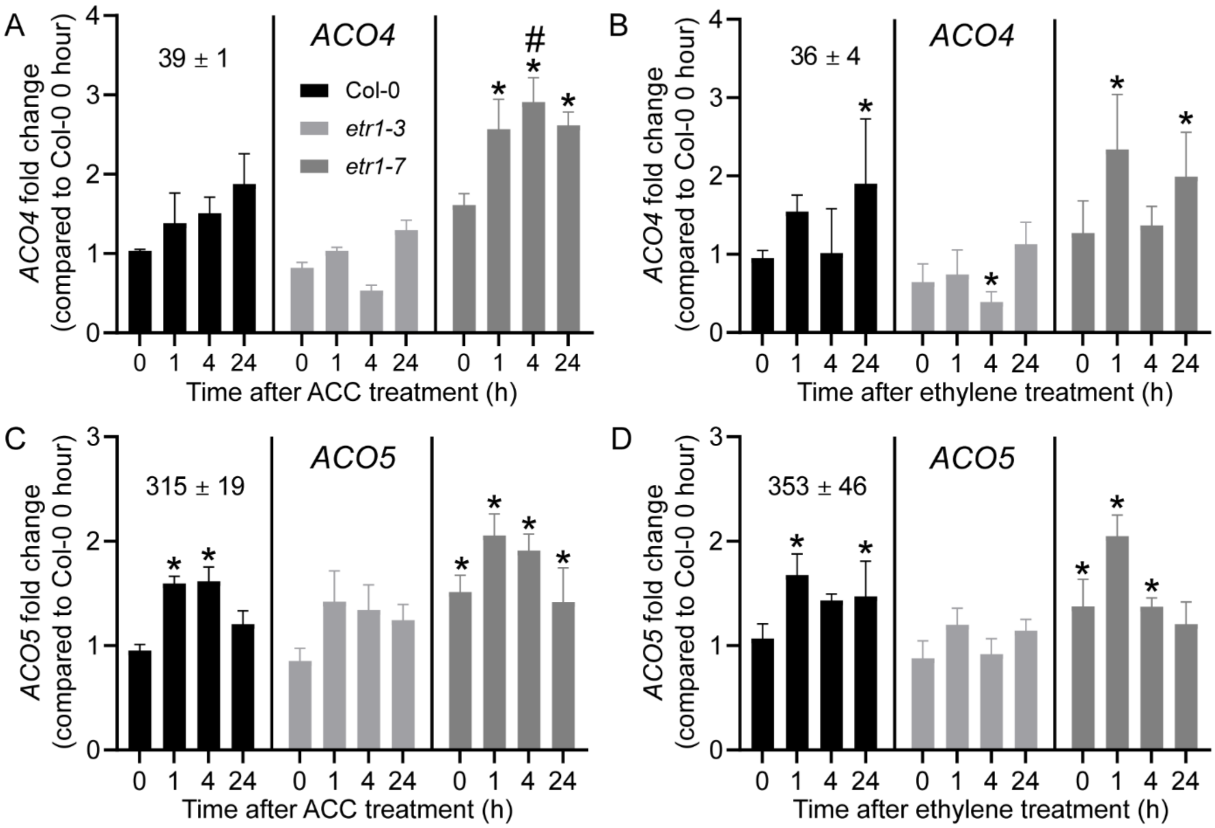

**Supplemental Figure 7. Transcript abundance of *ACO4* and *ACO5* in the *etr1* mutant alleles.** Transcript abundance (TPM) is reported relative to Col-0 time zero after ACC and ethylene treatment in our RNA-Seq dataset. Asterisks indicate statistical significance ($p < 0.01$) relative to untreated Col-0. Pound symbols indicate statistical significance ($p < 0.01$) compared to *etr1-7* time zero and are displayed only for the *etr1-7* data. Error bars represent SD.

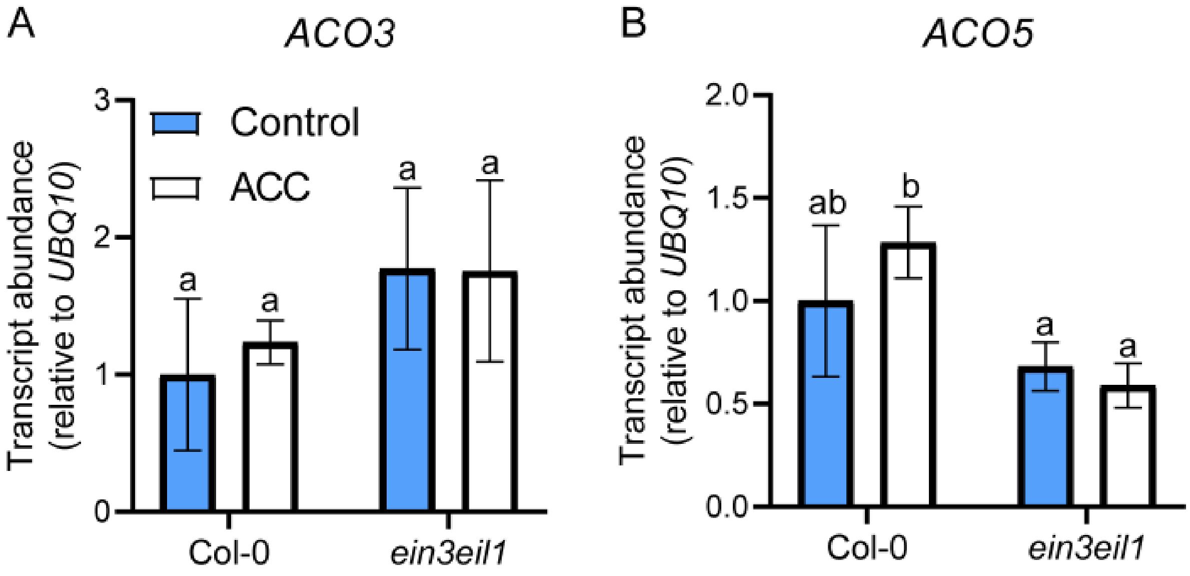

**Supplemental Figure 8. *ACO3* and *ACO5* transcript abundance is not significantly different in an *ein3/eil1* mutantEIN3/EIL1.** Transcript abundance in Col-0 and *ein3eil1* samples treated with and without 0.75 µM ACC for four hours. Error bars are SD. Statistical significance was defined as $p < 0.05$ according to Tukey's multiple comparisons test.

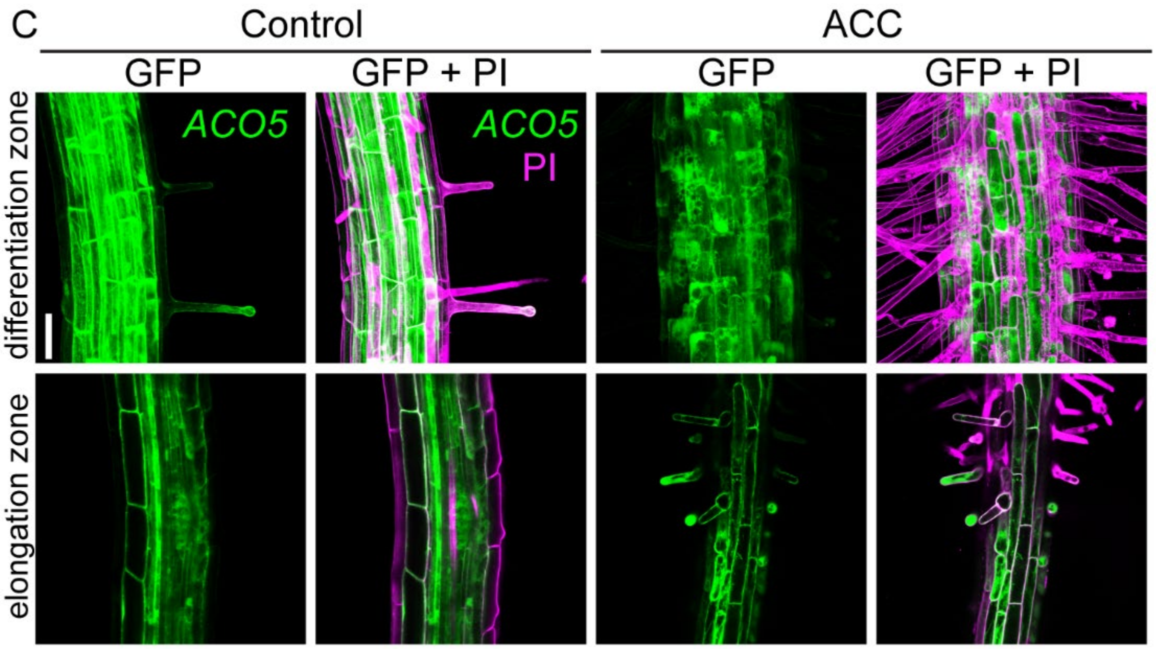

**Figure 9. *pACO5*::GFP-GUS fluorescence is expressed in root hairs.** Roots of 10-day-old *pACO5*::GFP-GUS seedlings in control or 0.75 µM ACC-treated for five days and stained with propidium iodide (PI; magenta) and imaged by laser scanning confocal microscopy. Scale bar= 150 µm.

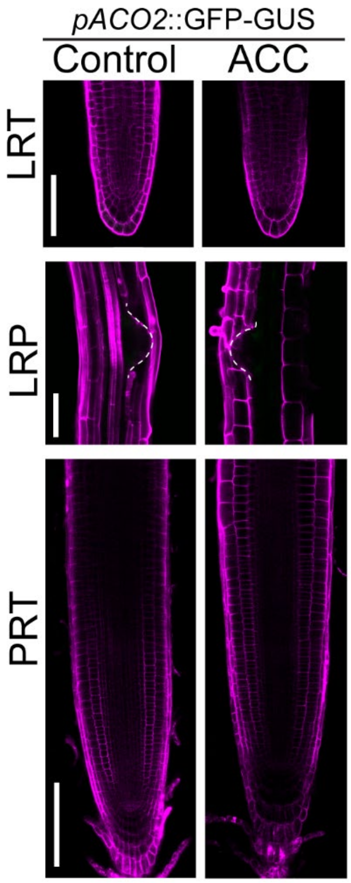

**Supplemental Figure 10.** ***ACO*2 was not regulated by ACC in tissues that control lateral root or primary root development.** 10-day-old *pACO2*::GFP-GUS seedlings treated with and without 0.75 µM ACC for five days. PRT, primary root tip. LRP, lateral root primordium. LRT, lateral root tip. Scale bar is 150 µm for PRT and LRT images and 100 µm for LRP images.

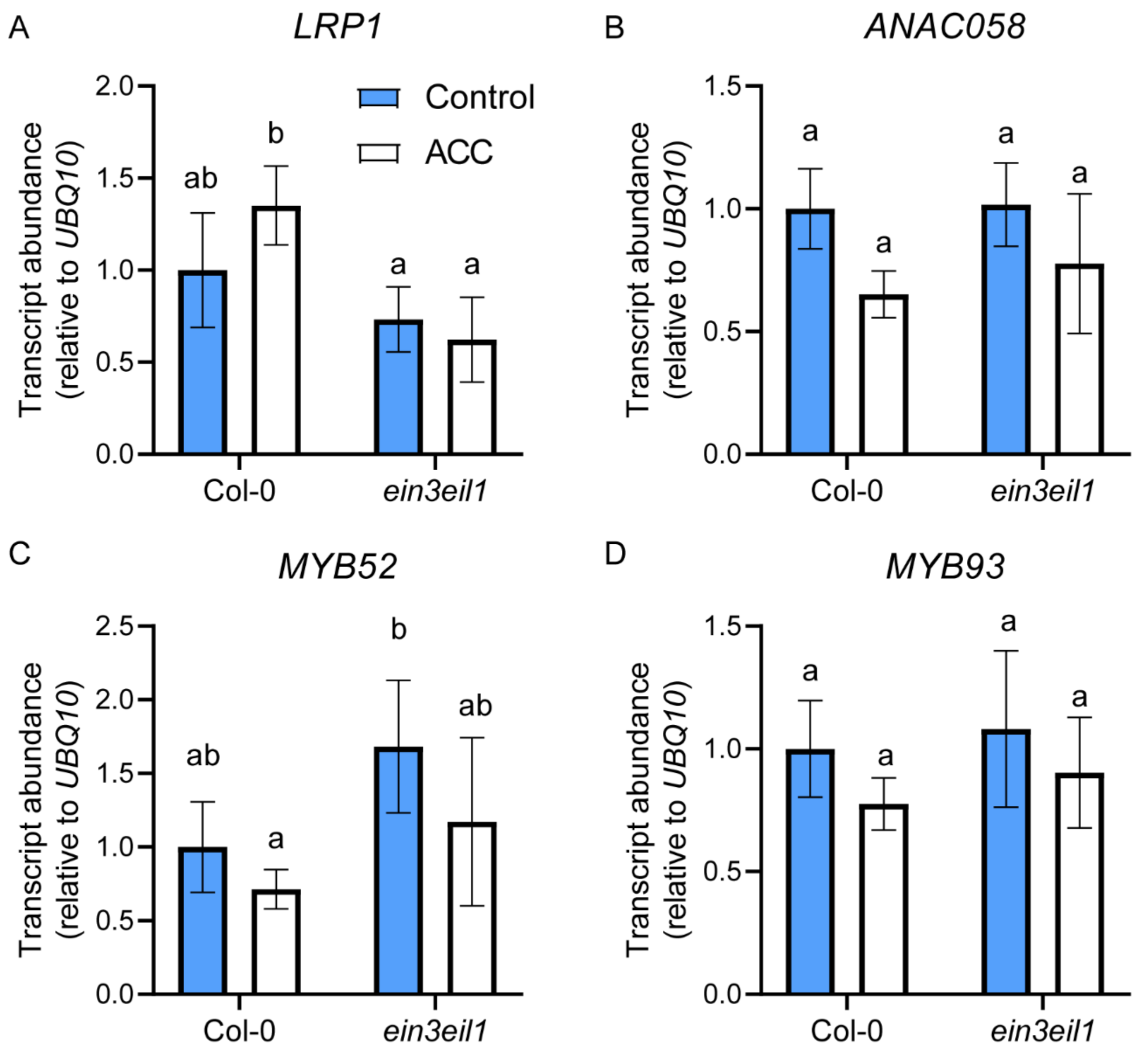

**Supplemental Figure 11. Several ETR1-dependent transcripts encoding TFs did not have significant changes in the *ein3eil1* mutant**. Transcript abundance in Col-0 and *ein3eil1* samples treated with and without 0.75 µM ACC for four hours. Error bars are SD. Statistical significance was defined as p-value < 0.05 according to Tukey's multiple comparisons test.

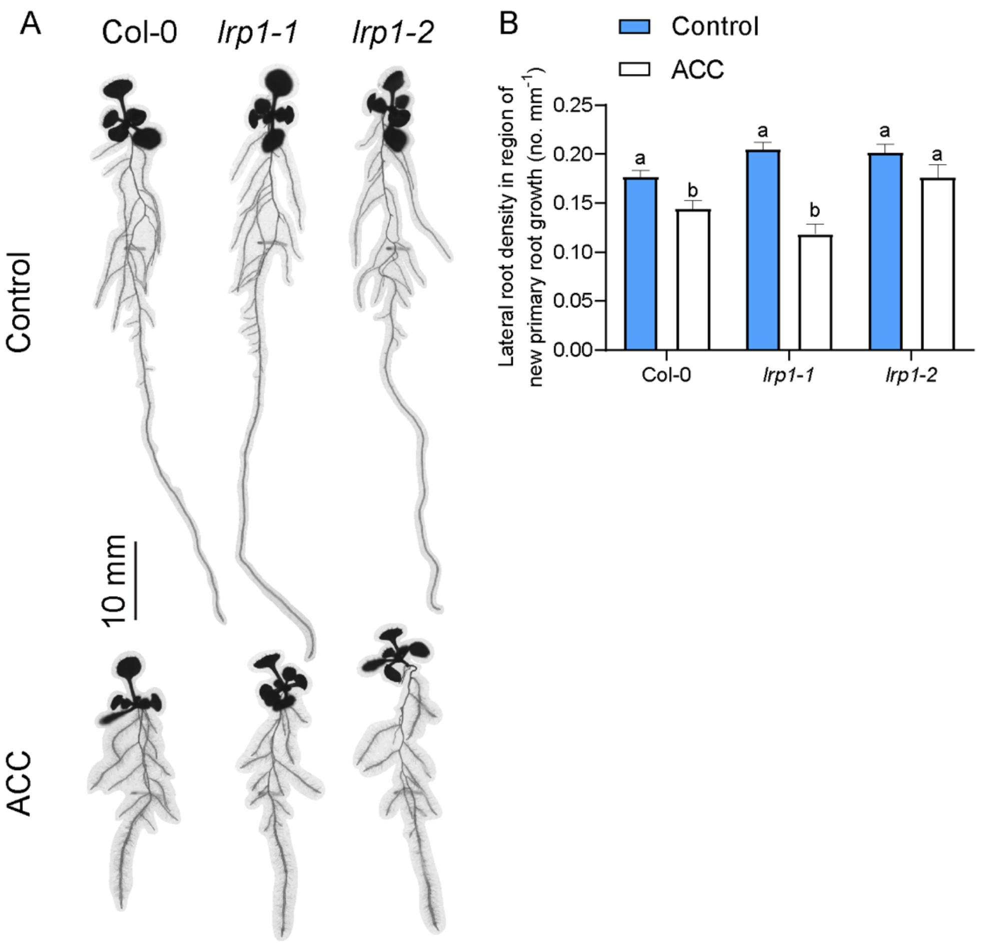

**Supplemental Figure 12. The *lrp1-1* and *lrp1-2* mutants had weak and not consistent root phenotypes.** (A) Representative images of 10-day-old Col-0, *lrp1-1*, and *lrp1-2* seedlings treated with and without 0.75 µM ACC for five days. Quantification of (B) the lateral root density in the region of new root growth in Col-0, *lrp1-1*, and *lrp1-2* in the presence and absence of 0.75 µM ACC after five days. Error bars are SD. Statistical significance is defined as p-value < 0.05 according to Tukey's multiple comparisons test.

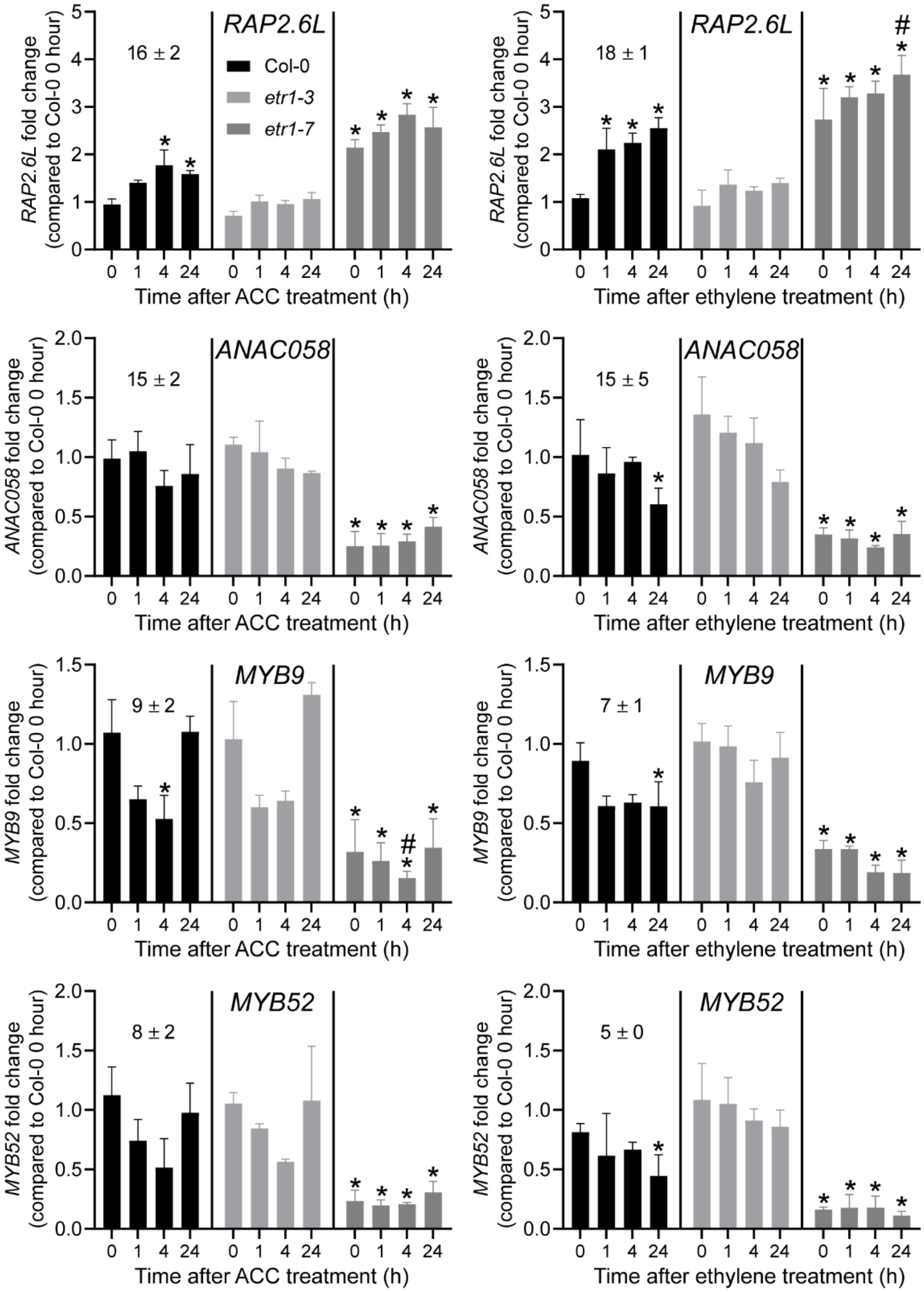

**Supplemental Figure 13. Transcript abundance of the four downregulated ETR1-dependent transcripts encoding TFs.** Transcript abundance (TPM) is reported relative to Col-0 time zero after ACC and ethylene treatment in our RNA Seq dataset. Asterisks indicate statistical significance (p-value < 0.01) relative to untreated Col-0. Pound symbols indicate statistical significance (p-value < 0.01) compared to *etr1-7* time zero and are displayed only for the *etr1-7* data. Error bars are SD.

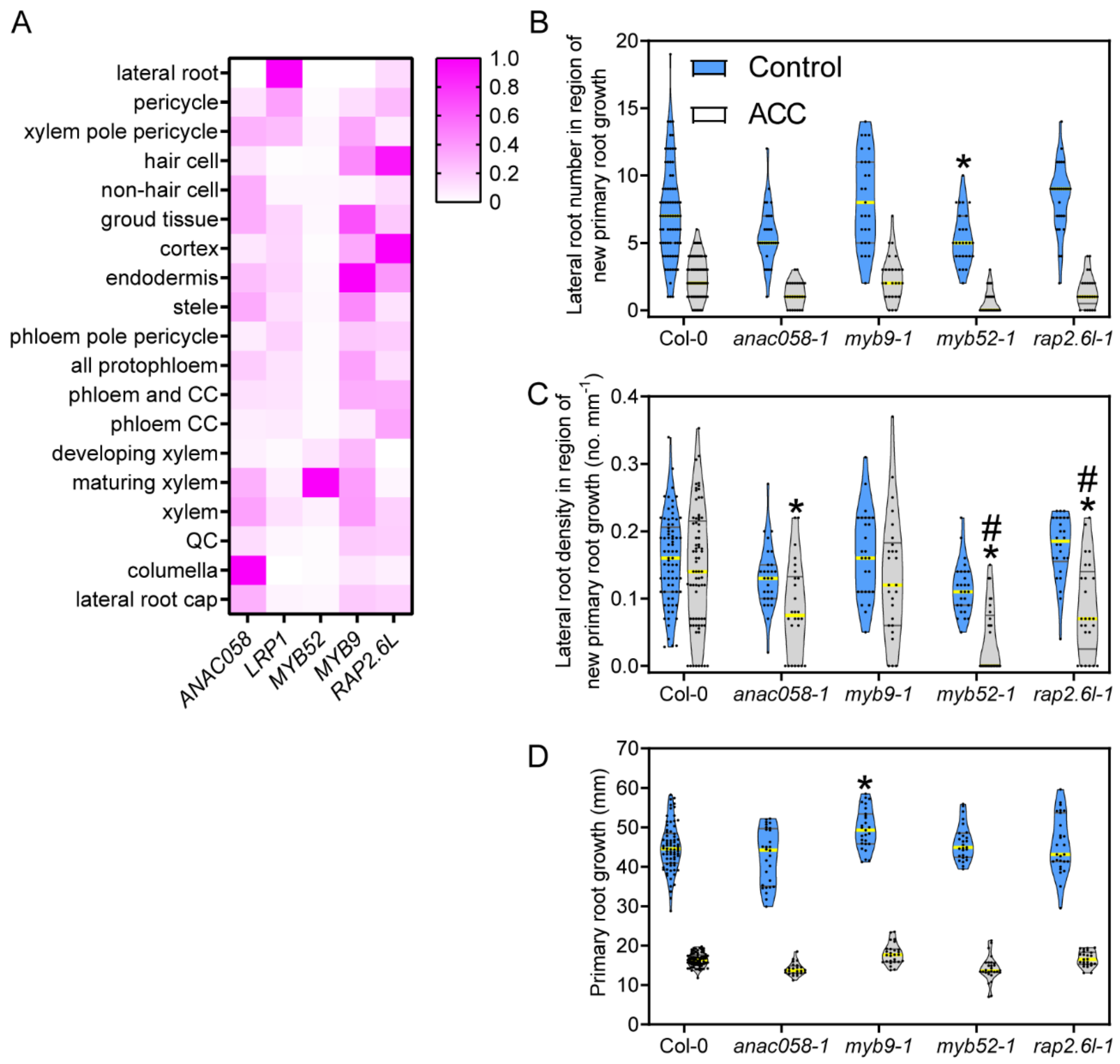

**Supplemental Figure 14. Mutants in several ETR1-dependent TFs have alterations in lateral root formation or primary root growth.** (A) Root cell type data reported as a heatmap (Brady et al. 2007). Data has been scaled and normalized to the tissue with the highest expression for each transcript. Violin plots of lateral root number in the region of new root growth (B), lateral root density in the region of new root growth (C), and primary root growth (D) for 10-day-old seedlings treated with and without 0.75 µM ACC for five days. Asterisks indicate significant differences from Col-0 under the same treatment conditions ($p$-value $< 0.05$). ACC effects within genotypes were significant for all genotypes in B and D (not shown on graphs). Pound symbols indicate significant differences within genotypes ($p$-value $<0.05$) for B. At least three biological replicates were performed. The median is illustrated in a thicker yellow line. The quartiles are illustrated in thinner gray lines.

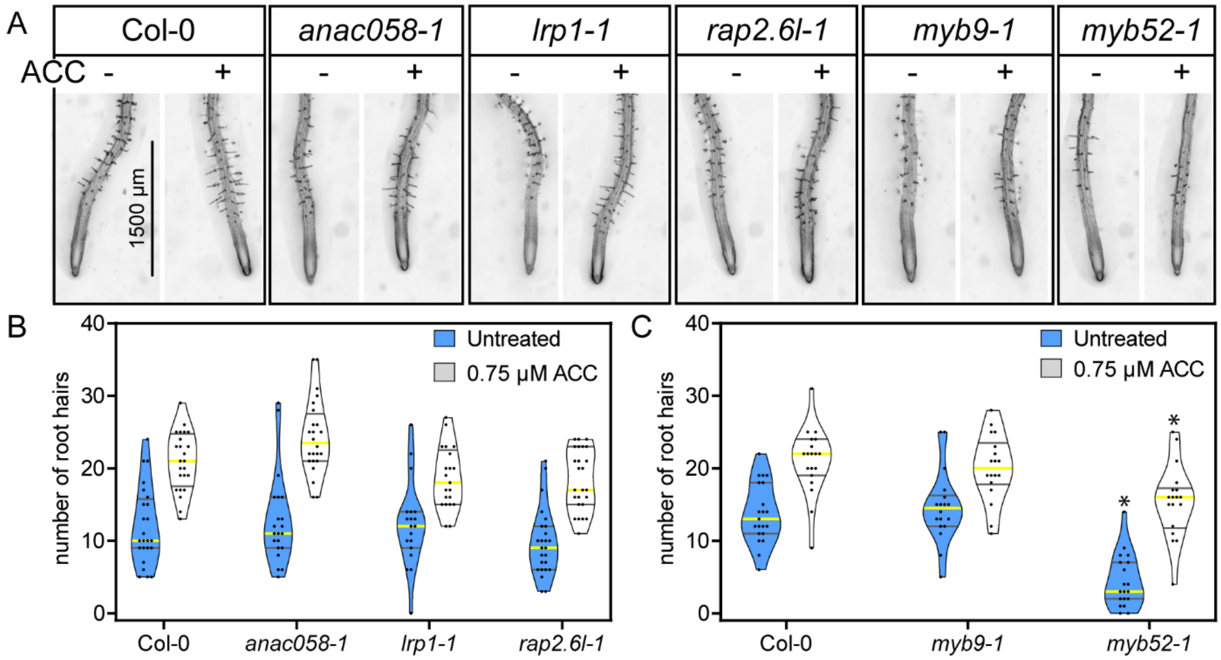

**Supplemental Figure 15. A mutant in the ETR1-dependent *MYB52* has altered root hair initiation.** (A) Representative images of five-day-old Col-0 and mutant seedlings treated with and without 0.75 µM ACC for four hours. Scale bar is 1,500 µm. (B,C) Violin plots of the number of root hairs in the region 0-1500 µm from the root tip in the absence and presence of 0.75 µM ACC. Asterisks indicate significant differences from Col-0 under the same condition, either untreated or with ACC ($p$-value $< 0.05$). ACC effects within genotypes were significant for all genotypes ($p$-value $< 0.05$). Three biological replicates were performed. The median is shown in a thicker yellow line. The quartiles are shown in thinner gray lines.

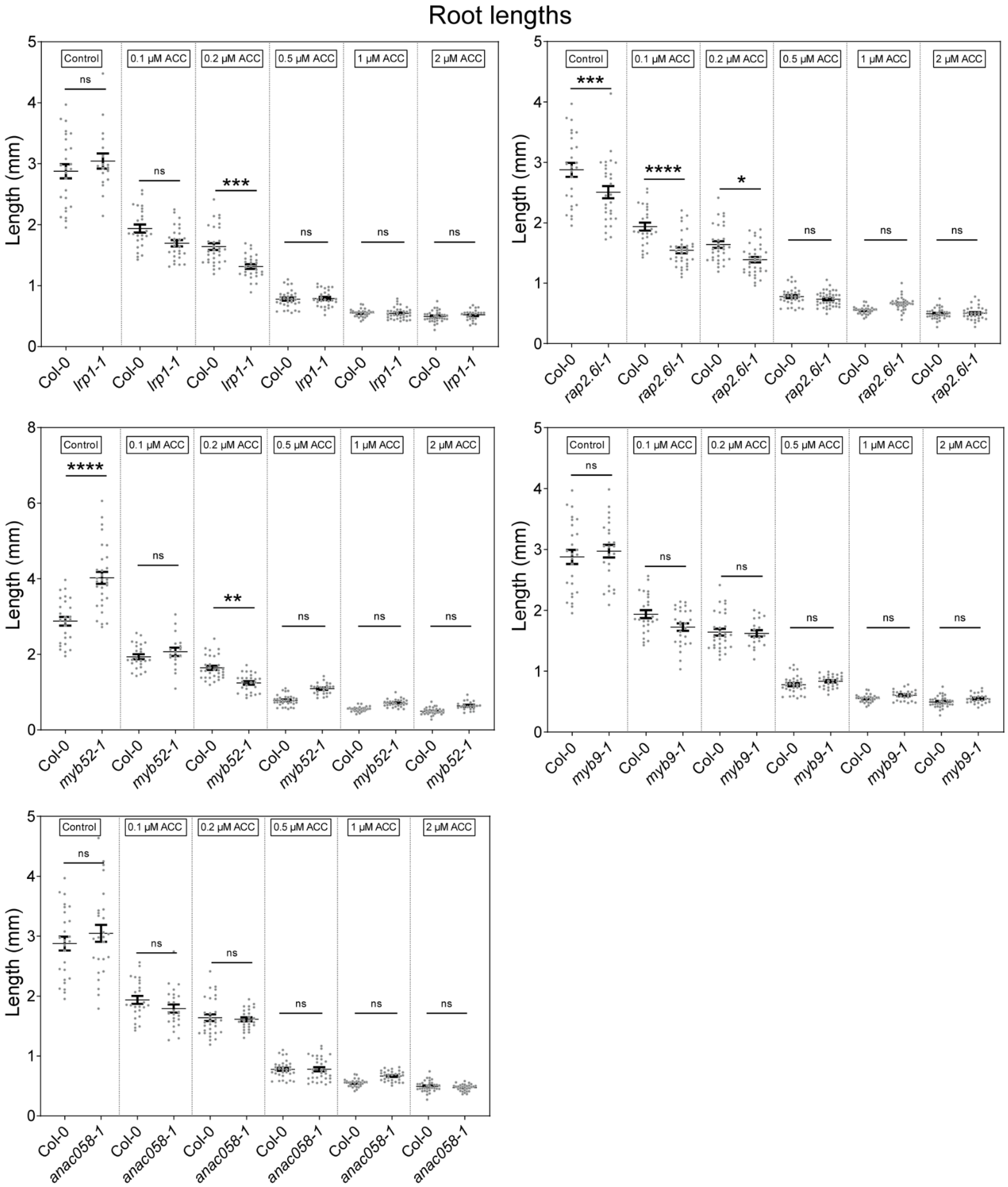

**Supplemental Figure 16. Triple response assay results for the candidate ETR1-dependent genes that function in root hair or lateral root formation.** Seedlings were incubated in the dark on control or ACC plates for four days before hypocotyl and root lengths were measured. Statistical significance was determined using a one-way ANOVA followed by a multiple comparisons test. * p-value < 0.05; ** p-value < 0.01; *** p-value < 0.001; **** p-value < 0.0001.

**Supplemental Table 1.** The number of differentially expressed genes between Wake Forest University- and University of Tennessee-prepared untreated samples for each genotype (with and without the untreated Col-0 outlier sample).

| Genotype | With outlier | Outlier removed |
|---|---|---|
| Col-0 | 693 | 34 |
| *etr1-3* | 137 | 121 |
| *etr1-7* | 997 | 763 |

**Supplemental Table 2.** Number of DE genes in response to ethylene and ACC based on two $log_2FC$ cutoffs. Note that the small number of genes that were regulated in opposite directions in the two treatments were not shown here for simplicity.

| Group of genes | 0.25 cutoff | 0.5 cutoff |
|---|---|---|
| Down in both | 860 | 504 |
| Down in ethylene | 2,461 | 1,493 |
| Down in ACC | 317 | 182 |
| Up in both | 620 | 457 |
| Up in ethylene | 2,563 | 1,657 |
| Up in ACC | 255 | 213 |

**Supplemental Table 3.** Gene Ontology (GO) enrichment results for genes transcriptionally regulated by ethylene and/or ACC in Col-0. Transcripts were separated based on if they responded to one or both treatments and if they were down- or up-regulated by the treatment(s). The number of genes with each GO annotation, the p-value and false discovery rate (FDR) for each annotation are reported.

| GO Term | # of genes | p-value | FDR |
|---|---|---|---|
| **Up in both treatments** | | | |
| regulation of transcription, DNA-dependent | 36 | 7.80E-09 | 5.00E-07 |
| response to chitin | 14 | 1.20E-08 | 5.70E-07 |
| regulation of gene expression | 50 | 1.10E-06 | 2.60E-05 |
| negative regulation of two-component signal transduction system (phosphorelay) | 5 | 1.70E-06 | 3.60E-05 |
| negative regulation of ethylene mediated signaling pathway | 5 | 1.70E-06 | 3.60E-05 |
| response to auxin stimulus | 16 | 1.30E-05 | 0.00025 |
| lipid localization | 5 | 2.10E-05 | 0.00039 |
| response to oxidative stress | 14 | 7.40E-05 | 0.0013 |
| response to osmotic stress | 14 | 0.00056 | 0.0095 |
| defense response to bacterium | 8 | 0.0017 | 0.027 |
| defense response, incompatible interaction | 7 | 0.0021 | 0.034 |
| response to external stimulus | 13 | 0.0025 | 0.04 |
| nitrogen compound metabolic process | 65 | 0.0027 | 0.042 |
| **Down in both treatments** | | | |
| lipid transport | 14 | 9.5E-08 | 0.000016 |
| hyperosmotic salinity response | 7 | 0.000014 | 0.0011 |
| response to cold | 16 | 0.000014 | 0.0011 |
| response to water deprivation | 13 | 0.00002 | 0.0013 |
| response to abscisic acid stimulus | 17 | 0.00002 | 0.0013 |
| response to nematode | 7 | 0.000066 | 0.0029 |
| phenylpropanoid biosynthetic process | 9 | 0.00016 | 0.0064 |

| | | | |
|---|---|---|---|
| response to oxidative stress | 14 | 0.0002 | 0.0076 |
| unidimensional cell growth | 10 | 0.00027 | 0.0094 |
| glycolysis | 5 | 0.0013 | 0.028 |
| terpenoid metabolic process | 7 | 0.0018 | 0.036 |
| **Up in ethylene only** | | | |
| chlorophyll biosynthetic process | 19 | 1.3E-11 | 1.7E-09 |
| cofactor biosynthetic process | 33 | 2.7E-10 | 2.8E-08 |
| chloroplast organization | 20 | 1.1E-09 | 1.1E-07 |
| response to high light intensity | 16 | 3.8E-08 | 3.4E-06 |
| lipid localization | 11 | 9.7E-08 | 8.1E-06 |
| photosynthesis, light harvesting | 11 | 1.8E-07 | 0.000014 |
| protein folding | 34 | 2.5E-07 | 0.000018 |
| response to red light | 17 | 2.9E-07 | 0.00002 |
| carbon fixation | 9 | 3.9E-07 | 0.000027 |
| photosynthetic electron transport in photosystem I | 10 | 5.5E-07 | 0.000034 |
| response to cold | 37 | 5.9E-07 | 0.000036 |
| response to blue light | 15 | 1.7E-06 | 0.000089 |
| embryonic development ending in seed dormancy | 45 | 1.9E-06 | 0.000097 |
| regulation of photosynthesis, light reaction | 8 | 2.4E-06 | 0.00012 |
| photosystem II assembly | 7 | 2.8E-06 | 0.00014 |
| response to far red light | 13 | 5.2E-06 | 0.00024 |
| protein import into chloroplast thylakoid membrane | 6 | 0.000011 | 0.00049 |
| response to other organism | 49 | 0.000043 | 0.0017 |
| response to hydrogen peroxide | 11 | 0.000058 | 0.0023 |
| defense response | 58 | 0.000068 | 0.0026 |
| thylakoid membrane organization | 7 | 0.00018 | 0.0062 |
| carotenoid biosynthetic process | 8 | 0.00024 | 0.0075 |
| positive regulation of development, heterochronic | 5 | 0.00025 | 0.008 |
| heat acclimation | 6 | 0.00045 | 0.013 |

| | | | |
|---|---|---|---|
| vitamin K biosynthetic process | 5 | 0.00049 | 0.014 |
| positive regulation of nitrogen compound metabolic process | 11 | 0.0005 | 0.014 |
| isopentenyl diphosphate biosynthetic process, mevalonate-independent pathway | 5 | 0.00066 | 0.018 |
| positive regulation of gene expression | 10 | 0.00082 | 0.022 |
| positive regulation of transcription | 10 | 0.00082 | 0.022 |
| positive regulation of cellular biosynthetic process | 11 | 0.00091 | 0.023 |
| xanthophyll metabolic process | 5 | 0.0014 | 0.034 |
| RNA metabolic process | 98 | 0.0017 | 0.039 |
| **Down in ethylene only** | | | |
| protein amino acid phosphorylation | 76 | 1.5E-08 | 4.7E-06 |
| oligopeptide transport | 17 | 1.5E-08 | 4.7E-06 |
| secondary cell wall biogenesis | 9 | 5.2E-07 | 0.000089 |
| glycoside metabolic process | 18 | 7.2E-07 | 0.0001 |
| plant-type cell wall organization | 15 | 2.2E-06 | 0.00026 |
| root hair cell differentiation | 10 | 0.000033 | 0.0021 |
| cell wall polysaccharide metabolic process | 6 | 0.000068 | 0.004 |
| transmembrane receptor protein tyrosine kinase signaling pathway | 16 | 0.00027 | 0.012 |
| glucosinolate metabolic process | 10 | 0.00034 | 0.014 |
| cell tip growth | 11 | 0.00055 | 0.02 |
| phenylpropanoid biosynthetic process | 15 | 0.00081 | 0.028 |
| lignin metabolic process | 8 | 0.00097 | 0.031 |
| multidrug transport | 10 | 0.0011 | 0.034 |
| isoprenoid biosynthetic process | 14 | 0.0015 | 0.046 |
| **Up in ACC only** | | | |
| regulation of biological quality | 12 | 0.00034 | 0.022 |
| defense response to bacterium | 6 | 0.0005 | 0.026 |

| | | | |
|---|---|---|---|
| protein amino acid phosphorylation | 14 | 0.00078 | 0.036 |
| transmembrane receptor protein tyrosine kinase signaling pathway | 5 | 0.0012 | 0.038 |
| response to inorganic substance | 7 | 0.00098 | 0.038 |
| response to hormone stimulus | 14 | 0.0011 | 0.038 |
| response to chitin | 5 | 0.0016 | 0.046 |
| **Down in ACC only** | | | |
| None significant | | | |
| **Up in either treatment** | | | |
| response to light intensity | 22 | 3.20E-07 | 2.00E-05 |
| cellular nitrogen compound biosynthetic process | 50 | 4.00E-06 | 0.00019 |
| fat-soluble vitamin biosynthetic process | 9 | 2.20E-05 | 0.00091 |
| toxin catabolic process | 13 | 7.80E-05 | 0.0027 |
| cellular response to chemical stimulus | 48 | 0.00033 | 0.011 |
| reductive pentose-phosphate cycle | 5 | 0.00034 | 0.011 |
| positive regulation of cellular metabolic process | 16 | 0.00037 | 0.012 |
| cellular response to stimulus | 76 | 0.00075 | 0.022 |
| quinone cofactor biosynthetic process | 8 | 0.00097 | 0.027 |
| response to iron ion | 5 | 0.0011 | 0.03 |
| cellular cation homeostasis | 14 | 0.0013 | 0.034 |
| **Down in either treatment** | | | |
| response to salt stress | 41 | 0.000091 | 0.0045 |
| lignin biosynthetic process | 7 | 0.00015 | 0.0065 |
| cell maturation | 11 | 0.00016 | 0.0069 |
| plant-type cell wall loosening | 10 | 0.00028 | 0.011 |
| sucrose biosynthetic process | 6 | 0.00084 | 0.028 |
| zinc ion transport | 6 | 0.00084 | 0.028 |
| fatty acid metabolic process | 26 | 0.0011 | 0.035 |

| | | | |
|---|---|---|---|
| glucosinolate biosynthetic process | 9 | 0.0012 | 0.039 |
| pentacyclic triterpenoid biosynthetic process | 5 | 0.0016 | 0.049 |
| alcohol metabolic process | 29 | 0.0016 | 0.049 |
| abscisic acid mediated signaling pathway | 8 | 0.18 | 1 |
| oligopeptide transporter activity | 5 | 0.011 | 0.14 |

**Supplemental Table 4.** GO annotations enriched in genes which respond to ethylene or ACC via ETR1-dependent and -independent pathways. The number of genes with each GO annotation, the p-value and false discovery rate (FDR) for each annotation are reported.

| GO Term | # of genes | p-value | FDR |
|---|---|---|---|
| **ETR1 dependent (all)** | | | |
| oligopeptide transport | 7 | 0.000092 | 0.0097 |
| lipid transport | 10 | 0.00019 | 0.011 |
| post-embryonic development | 24 | 0.00013 | 0.011 |
| response to water deprivation | 12 | 0.00018 | 0.011 |
| regulation of transcription, DNA-dependent | 30 | 0.00014 | 0.011 |
| response to abscisic acid stimulus | 15 | 0.00053 | 0.021 |
| ethylene mediated signaling pathway | 6 | 0.00095 | 0.03 |
| fatty acid metabolic process | 10 | 0.002 | 0.048 |
| **ETR1 independent (all)** | | | |
| response to heat | 21 | 1.60E-11 | 8.30E-09 |
| response to chitin | 17 | 1.00E-08 | 3.20E-06 |
| response to hydrogen peroxide | 8 | 1.20E-05 | 0.0013 |
| response to high light intensity | 8 | 2.00E-05 | 0.0018 |
| root hair cell differentiation | 7 | 3.70E-05 | 0.0027 |
| unidimensional cell growth | 12 | 0.00025 | 0.013 |
| protein folding | 14 | 0.00076 | 0.036 |
| inorganic anion transport | 5 | 0.00087 | 0.04 |
| nucleotide-sugar metabolic process | 5 | 0.00097 | 0.043 |
| cellular response to nutrient levels | 7 | 0.0011 | 0.045 |
| lipid metabolic process | 29 | 0.0011 | 0.045 |
| plant-type cell wall loosening | 5 | 0.0012 | 0.048 |

| ETR1 complex (all) | | | |
|---|---|---|---|
| response to light stimulus | 180 | 1.4E-08 | 0.000004 |
| response to chitin | 64 | 1.1E-07 | 0.000026 |
| phenylpropanoid biosynthetic process | 60 | 2.4E-07 | 0.000051 |
| toxin catabolic process | 33 | 2.5E-07 | 0.000051 |
| protein amino acid phosphorylation | 244 | 2.6E-06 | 0.00044 |
| oligopeptide transport | 35 | 3.1E-06 | 0.00048 |
| response to wounding | 71 | 2.9E-06 | 0.00048 |
| glucosinolate biosynthetic process | 25 | 8.9E-06 | 0.0011 |
| plant-type cell wall organization | 37 | 8.7E-06 | 0.0011 |
| monocarboxylic acid metabolic process | 120 | 0.000009 | 0.0011 |
| response to nematode | 34 | 0.000011 | 0.0013 |
| response to salt stress | 107 | 0.000033 | 0.0037 |
| post-embryonic morphogenesis | 21 | 0.000054 | 0.0055 |
| photosynthetic electron transport chain | 24 | 0.000088 | 0.0086 |
| multidrug transport | 32 | 0.000091 | 0.0088 |
| response to abscisic acid stimulus | 107 | 0.000094 | 0.009 |
| response to metal ion | 74 | 0.00011 | 0.0097 |
| intracellular signaling cascade | 169 | 0.00011 | 0.0097 |
| cell wall modification | 45 | 0.00013 | 0.012 |
| response to cold | 94 | 0.00017 | 0.015 |
| disaccharide biosynthetic process | 21 | 0.00017 | 0.015 |
| protein complex assembly | 47 | 0.00021 | 0.017 |
| heterocycle metabolic process | 123 | 0.00022 | 0.018 |
| defense response to fungus | 40 | 0.00025 | 0.019 |
| terpenoid biosynthetic process | 36 | 0.00029 | 0.022 |
| cellular carbohydrate catabolic process | 44 | 0.00031 | 0.023 |
| regulation of transcription | 400 | 0.00039 | 0.028 |

| | | | |
|---|---|---|---|
| response to ethylene stimulus | 61 | 0.00055 | 0.034 |
| cellular glucan metabolic process | 33 | 0.00059 | 0.036 |
| cation transport | 97 | 0.00062 | 0.037 |
| two-component signal transduction system (phosphorelay) | 34 | 0.00071 | 0.042 |
| cell wall macromolecule metabolic process | 21 | 0.00074 | 0.044 |
| chemical homeostasis | 45 | 0.00078 | 0.045 |
| response to water deprivation | 67 | 0.00085 | 0.049 |
| transition metal ion transmembrane transporter activity | 16 | 0.022 | 0.43 |

**Supplemental Table 5. Genotyping primers used in this study.** Sequences are written 5' to 3'. L, left border primer. R, right border primer.

| Genome Locus | Allele Name | T-DNA Insertion Line | Location of Insertion | Primer Sequences |
|---|---|---|---|---|
| AT3G18400 | *anac058-1* | SALK_049205C | 3' UTR | L:GTTCTGGCAGCAATTGTGATC<br>R:CAACTCTTTGTTGCTCAAGGC |
| AT5G12330 | *lrp1-1* | SAIL_402_G06/ CS873836 | Promoter | L:TGTATGGCCAGTAAGCAATCC<br>R:ACGTCGGTAGATTATGCCTCC |
| AT5G12330 | *lrp1-2* | SALK_201247C | Exon | L:TGATAAGTGCATGAGAAAATGG<br>R:ACACACCACCTCAAAGTTTCG |
| AT5G16770 | *myb9-1* | SALK_149765C | Promoter | L:TGGAATCGCAGTTTTTCTAGC<br>R:CGTGCATAAATATTTGCATGTG |
| AT1G17950 | *myb52-1* | SALK_138624C | Exon | L:AAAAGGATGTTCATTTGGTGG<br>R:TGCAAGTAAATGAGTAATGGTGC |
| AT5G13330 | *rap2.6l-1* | SALK_051006C | 5' UTR | L:AAAAGGTGGTACATACAGGCG<br>R:TCGGTGGTTCAATTCAAAAAC |

**Supplemental Table 6. RT-qPCR primers used in this study.** Sequences are written 5' to 3'. F, forward primer. R, reverse primer.

| Genome Locus | Gene Symbol | Primer Sequences |
|---|---|---|
| AT2G19590 | *ACO1* | F: TGGTCCAAAAGGTCCAGCTTT<br>R: ATTCCCCCAGCATCCGTATG |
| AT1G62380 | *ACO2* | F: TCTACGTTCGTCACCTCCCT<br>R: AGTCTTTCATGGCCGTCCTG |
| AT1G12010 | *ACO3* | F: ACAACGGGTCCAACTTTTGC<br>R: TGTGAGCCCTAAGCCCTTTG |
| AT1G05010 | *ACO4* | F: CTACCTCAAGCACCTTCCCG<br>R: GCACAGCAGATCCAGTAGCT |
| AT1G77330 | *ACO5* | F: AGTCAATGGCCTTCGAGCTC<br>R: GATCCACTCGCCGTCTTTCA |
| AT3G18400 | *ANAC058* | F: ACAACCTTTCAACCCCACGA<br>R: TTTGCTGCCGTGCTCTTTTC |
| AT5G12330 | *LRP1* | F: CTGTGGCTACAACGAGGAGG<br>R: ACGGAAGCTGAGCCTAAACC |
| AT5G16770 | *MYB9* | F: ACAAGTGGTCGTCGATAGCC<br>R: GGCTATAAGCTGCGGGAGAG |
| AT1G17950 | *MYB52* | F: CCGGTCGAACTGATAACGCT<br>R: TAGCGGTTGTGGTTGCCAAT |
| AT1G34670 | *MYB93* | F: ATCAACCCACCCAACCACTC<br>R: GTCACCTTGCGATGGTAGCT |
| AT5G13330 | *RAP2.6L* | F: CAAAGAAAGCAGCCCGTGTC<br>R: GTTGTGGTGGTAGTAGGGCC |
| AT5G43175 | *RSL5* | F: GCAATTTGGTTACCGAGCCT<br>R: ATCCCTCTGTTGGCTTTCGC |
